# Enhancing gene transfer to renal tubules and podocytes by context-dependent selection of AAV capsids

**DOI:** 10.1101/2023.07.28.548760

**Authors:** Taisuke Furusho, Kei Adachi, Mia Galbraith-Liss, Anusha Sairavi, Ranjan Das, Hiroyuki Nakai

## Abstract

Despite recent remarkable advancements in adeno-associated virus (AAV) vector technologies, effective gene delivery to the kidney remains a significant challenge. Here we show that AAV vector transduction in proximal tubules and podocytes, the crucial targets for renal gene therapy, can be enhanced remarkably through a meticulous selection of both AAV capsids and route of administration, tailored to the condition of the kidney. In this study, we performed a side-by-side comparison of 47 AAV capsids using AAV Barcode-Seq and identified six AAV capsids, including AAV-KP1, that exhibit remarkable enhancement of renal transduction in mice when delivered locally via the renal vein or the renal pelvis. Individual capsid validation analyses revealed that local delivery of AAV-KP1, but not AAV9, enables remarkably enhanced proximal tubule transduction while minimizing off-target liver transduction. In a mouse model of chronic kidney disease, intravenous administration of AAV9, not AAV-KP1, showed efficient renal tubule and podocyte transduction, which was not observed in the control wild-type mice. We also provide evidence that these contrasting observations between AAV-KP1 and AAV9 are attributed to their distinct pharmacokinetic profiles. Thus, this study highlights the importance of context-dependent capsid selection and engineering for successful renal gene therapy.

## INTRODUCTION

Adeno-associated virus (AAV) vector-mediated gene therapy has shown promise in treating genetic diseases, and commercial AAV gene therapy products have been approved by regulatory agencies and used in the clinic to treat spinal muscular atrophy, Leber congenital amaurosis, hemophilia A, hemophilia B, aromatic L-amino acid decarboxylase (AADC) deficiency and Duchenne muscular atrophy.^1–5^ With recent remarkable progress in the understanding of genetic etiology of kidney diseases, the success of AAV vector-mediated gene therapy has brought a significant hope to patients who suffer from genetic kidney diseases that account for a significant fraction of both pediatric and adult patients with chronic kidney disease (CKD).^6–8^ However, in contrast to other organs, gene delivery to the kidney has been challenging and hampered by its low transduction efficiency.^9^ Intravenous (IV) administration of common AAV serotype vectors only achieves mesangial cell and interstitial cell transduction in the kidney, and the current method cannot target effectively clinically relevant cell types including tubular epithelial cells and podocytes.^10, 11^ In addition, although recent advancement of capsid engineering technology generated many AAV capsids with novel phenotypes,^1, 2^ they have not been characterized in detail in the context of renal gene transfer. Thus, the development of AAV vector-mediated renal gene therapy has been impeded so far.

Local AAV vector administration has a potential to enhance target tissue transduction and prevent off-target effect. For renal gene transfer, injection into the renal circulation such as retrograde renal vein (RV) injection and injection into the urinary tract such as retrograde renal pelvis (RP) injection are the two common local routes of administration.^12–16^ To date, however, there has been no study that compared renal transduction efficiencies of various AAV serotypes and capsid mutants by RV and RP injections in a head-to-head setting. Thus, administration route-dependent differences in renal transduction profiles among different AAV capsids remain not well understood. Regarding off-target effects, minimizing vector dissemination to off-target organs is important to prevent serious side effects including hepatotoxicity, genotoxicity, thrombotic microangiopathy and dorsal root ganglia (DRG) toxicity.^1, 17^ However, recent studies have shown substantial off-target liver transduction of AAV8 and AAV9 following RV^12^ and RP^13^ injections, indicating that these commonly used AAV capsids are not ideal for local injection. Since off-target organ transduction profiles of various AAV capsid-derived vectors after RV or RP injection have not been assessed before, it has yet to be determined which AAV capsids are suitable for local kidney injection.

CKD is characterized by reduced glomerular filtration and proteinuria. In addition to proteinuria due to increased permeability of the glomerular filtration barrier (glomerular capillary), increased peritubular capillary permeability is also a common feature of CKD.^18^ As vascular endothelial cells are the major barrier for AAV transduction,^19^ different renal transduction profiles are expected between healthy and CKD kidneys. Moreover, assessment of renal transduction in CKD is imperative since AAV vector-mediated renal gene therapy may need to be tailored to patients with CKD. Nevertheless, renal transduction profiles of different AAV capsids in CKD have never been determined so far.

In the present study, we comprehensively assessed renal transduction efficiencies of currently available AAV capsids by IV, RV and RP injections using AAV Barcode-Seq (Nakai, H., Huang, S., Adachi, K. US11,459,558. October 4, 2022.).^20^ As a result, we identified six AAV capsids (AAV-KP1, AAV-KP2, AAV-KP3, AAV-DJ, AAV2G9, AAV2.7m8) that show remarkable enhancement of renal transduction and transduce the kidney much more efficiently than AAV9 following RV and RP injections despite showing no advantages when delivered via IV. Detailed analysis revealed that local injection of AAV-KP1 enables gene transfer to proximal tubules while minimizing the off-target liver transduction. On the other hand, there was no advantage of local injection of AAV9 whose renal and hepatic transduction remained the same regardless of administration routes. Finally, this study demonstrated for the first time that renal transduction profile is substantially different between healthy and CKD kidneys. Especially, AAV9 efficiently transduces proximal tubules and podocytes in CKD kidneys following IV administration. These observations not only help us design renal gene delivery approaches that enhance transduction in target cells but also emphasize the importance of tailored AAV vector selection based on the administration route and the host condition.

## RESULTS

### AAV Barcode-Seq identified AAV capsids that outperform AAV9 in renal transduction by local injection

We first investigated which AAV capsids transduce the kidney most efficiently by IV, RV and RP injections in mice. To this end, we employed AAV Barcode-Seq (Nakai, H., Huang, S., Adachi, K. US11,459,558. October 4, 2022.) ^20^ and determined renal transduction profiles of a total of 47 different AAV capsids in mice following IV, RV or RP administration. In this approach, AAV capsids package the AAV-CAG-BC vector genomes whose barcodes (BCs) are unique to each capsid (**Figure 1A**). The AAV-CAG-BC genomes carry the ubiquitous CAG promoter that drives barcode expression as mRNA. This design allows the assessment of relative transduction efficiency at both DNA and RNA levels (*i.e.*, DNA/RNA Barcode-Seq). We produced an AAV barcode library that contained 47 AAV capsids (11 naturally occurring serotypes and 36 capsid-engineered mutants, **Table S1**) and injected the library via IV (2x10^13^ vg/kg), RV (3x10^11^ vg/mouse) or RP (3x10^11^ vg/mouse) into eight-week-old C57BL/6J male mice (n=3 for IV and RP, n=4 for RV). RV and RP injections were performed with the blockage of the local renal blood flow for 15 min based on the methods reported previously^12, 16^ to increase the vector exposure. For RP injection, urinary flow was also blocked, and the renal pelvis was slowly filled with the AAV vector solution over 1 min. Six weeks post-injection, transduction efficiency of each AAV capsid in the kidney relative to that of AAV9 was determined by AAV DNA/RNA Barcode-Seq. As shown in **Figure 1B** (Note: Transduction efficiencies of all AAV capsids are shown in **Figure S1**), we found that (a) AAV9 capsid transduced the kidney most efficiently by IV; (b) by RV and RP injections, the following six AAV capsids, AAV-KP1, AAV-KP2, AAV-KP3,^21^ AAV-DJ,^22^ AAV2G9^23^ and AAV2.7m8^24^ showed significantly higher renal transduction than AAV9 at the level of vector genome mRNA transcripts (A one-way ANOVA followed by Tukey’s post hoc test was used. For RV injection, adjusted p=4.0x10^-13^ (AAV-KP1), 1.7x10^-10^ (AAV-KP2), 2.9x10^-13^ (AAV-KP3), 3.7x10^-10^ (AAV-DJ), 3.4x10^-13^ (AAV-2G9), 1.7x10^-2^ (AAV2.7m8). For RP injection, adjusted p=4.8x10^-10^ (AAV-KP1), 4.8x10^-10^ (AAV-KP2), 4.8x10^-10^ (AAV-KP3), 4.8x10^-10^ (AAV-DJ), 4.8x10^-10^ (AAV-2G9), 8.5x10^-4^ (AAV2.7m8).); (c) there was a DNA/RNA correlation for these six AAV capsids; and (d) AAV3, AAV-LK03 and AAVShH10 showed significantly higher renal gene transfer than AAV9 at the vector genome DNA level (A one-way ANOVA followed by Tukey’s post hoc test was used. For RV injection, adjusted p=3.0x10^-14^ (AAV3), 1.0x10^-15^ (AAV-LK03), 1.0x10^-15^ (AAVShH10). For RP injection, adjusted p=4.8x10^-10^ (AAV3), 4.8x10^-10^ (AAV-LK03), 4.8x10^-13^ (AAVShH10).) but the vector genomes delivered by these three capsids were transcriptionally less active than those delivered by AAV9 with statistical significance (A one-way ANOVA followed by Tukey’s post hoc test was used. For RV injection, adjusted p=1.6x10^-11^ (AAV3), 1.3x10^-4^ (AAV-LK03), 1.7x10^-4^ (AAVShH10). For RP injection, adjusted p=4.8x10^-10^ (AAV3), 1.3x10^-9^ (AAV-LK03), 2.1x10^-9^ (AAVShH10).), showing a discordance between the AAV vector genome DNA levels and the vector genome mRNA transcript levels. Since this phenotype of AAV-LK03 was previously reported in the liver^25^ and the discordance between vector genome DNA and RNA transcript levels was also observed with AAV3, AAV-LK03 and AAVshH10 in IV injection (**Figure S1**), this phenomenon is most likely not related to the route of vector administration. Importantly, systemic and local vector administrations showed distinct profiles in renal transduction. That is, the observed enhancement in renal transduction with the six AAV capsids we identified was specific to local administration, emphasizing the critical need to carefully evaluate administration route-dependent attributes of AAV capsids when designing gene therapy approaches for the treatment of kidney diseases.

**Figure 1.**
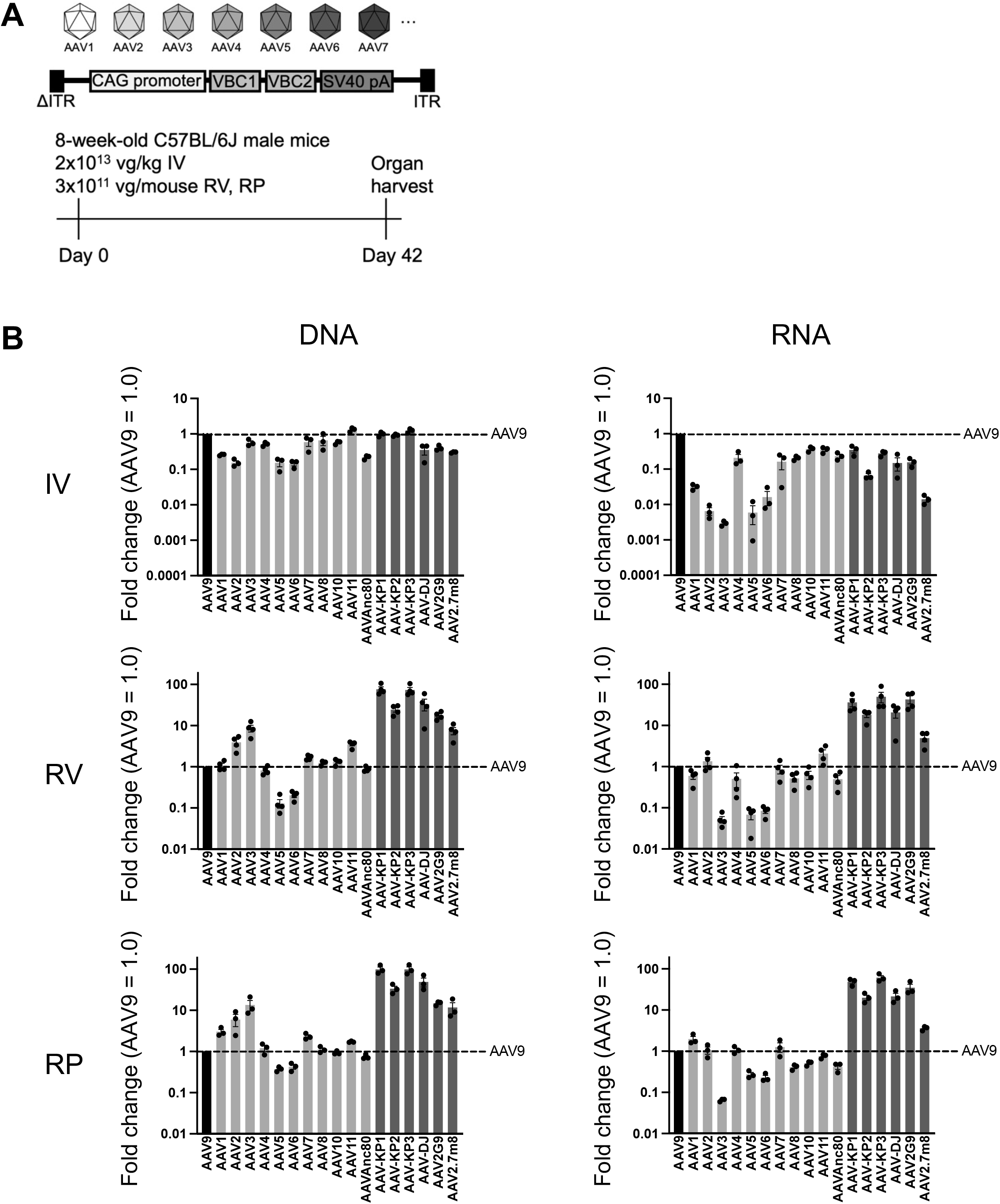
AAV Barcode-Seq analysis of AAV capsids for renal transduction following intravenous (IV), renal vein (RV) and renal pelvis (RP) injections. **(A)** The experimental design. Eight-week-old C57BL/6J male mice were injected with an AAV-CAG-Barcode library that contained 47 AAV capsids via IV (2x10^13^ vg/kg), RV (3x10^11^ vg/mouse) or RP (3x10^11^ vg/mouse) (n=3 for IV and RP, n=4 for RV). Kidneys were harvested 6 weeks post-injection for the AAV Barcode-Seq analysis^20^ (Nakai, H., Huang, S., Adachi, K. US11,459,558. October 4, 2022.). **(B)** Relative quantities of each AAV capsid-derived vector genomes (DNA) and vector genome transcripts (RNA) in the kidney were determined by AAV Barcode-Seq and expressed as fold changes. The light and dark bars represent common AAV serotypes and the six AAV capsids showing significantly enhancement RNA transduction compared to AAV9, respectively. Error bars represent SEM. A one-way ANOVA followed by Tukey’s post hoc test was used for statistical assessment of the data. Adjusted p<0.05 was considered as statistical significance. For the complete dataset of the 47 AAV capsids, refer to Figure S1.

### Local injection of AAV-KP1 enables effective proximal tubule cell transduction

To validate the AAV Barcode-Seq data and characterize the cell type tropism, we individually packaged the AAV-CAG-tdTomato genome with AAV9 and AAV-KP1 capsids and produced AAV9-CAG-tdTomato and AAV-KP1-CAG-tdTomato vectors. Each of these two vectors was injected into eight-week-old C57BL/6J male mice by IV, RV or RP at a dose of 3x10^11^ vg/mouse (n=4 per group, **Figure 2A**). We selected AAV-KP1 from the six AAV capsids showing the enhancement as it was one of the most efficient capsids for kidney transduction by both RV and RP based on the AAV Barcode-Seq data. Two weeks post-injection, vector genome copy numbers in the kidney were determined by DNA qPCR and renal transduction was assessed by fluorescence microscopy. Quantification of vector genomes in the kidney showed that, while vector genome copy numbers were comparable between the AAV9 and AAV-KP1-injected mice in the IV group, there were 35 times and 23 times more vector genomes in the AAV-KP1-injected mice than the AAV9-injected mice following RV and RP injections, respectively (**Figure 2B**). Vector genome copy numbers in the AAV9-injected animals were not different between the IV, RV and RP groups, demonstrating that there is no advantage of local administration over systemic administration when AAV9 capsid is used. In consistent with the vector genome copy number data, the histological assessment also showed substantially enhanced transduction in the kidney with AAV-KP1 following RV and RP injections compared to IV injection (**Figure 2C**). Interestingly, enhanced transduction of AAV-KP1 was mainly observed in the cortex, specifically the proximal tubules by co-staining with Lotus tetragonolobus lectin (LTL). The proximal tubular cell transduction was 11% and 6.5% by RV and RP injections of AAV-KP1, respectively, whereas AAV-KP1 transduced only mesangial cells in the glomeruli by IV injection (**Figure 3A, 3C**). On the other hand, the cell tropism of AAV9 was not altered regardless of the administration routes, and showed primarily mesangial cell transduction. Co-staining with a podocyte marker, Wilms tumor 1 (WT1), demonstrated no appreciable transduction in podocytes with AAV9 or AAV-KP1 regardless of the administration routes (**Figure 3B**).

**Figure 2.**
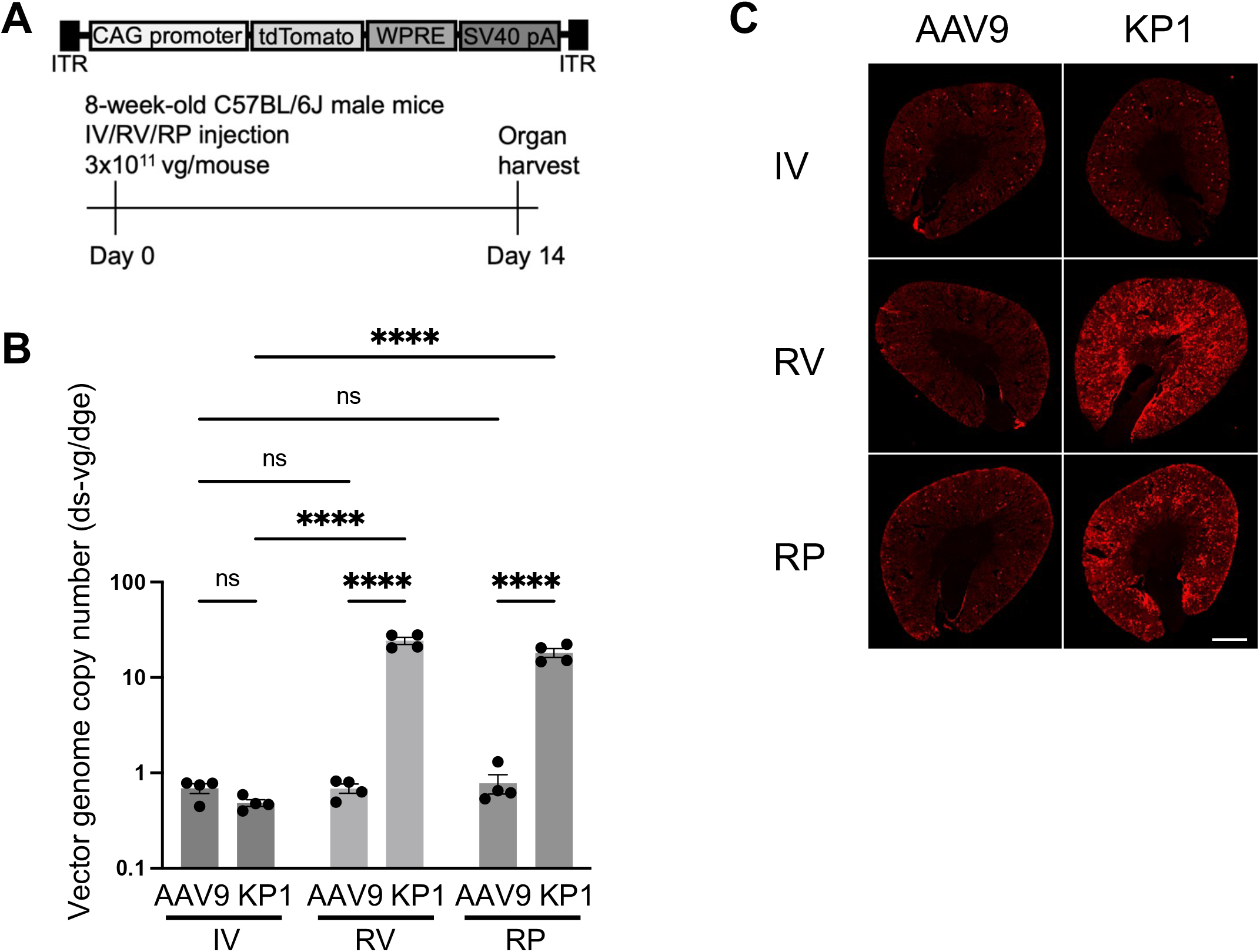
Enhanced renal transduction with AAV-KP1 by RV or RP injection. **(A)** The experimental design. Eight-week-old C57BL/6J male mice were injected with AAV9-CAG-tdTomato or AAV-KP1-CAG-tdTomato via RV or RP at a dose of 3x10^11^ vg/mouse (n=4 per group). Kidneys were harvested 2 weeks post-injection for downstream analyses. **(B)** AAV vector genome copy numbers in the kidneys determined by qPCR. Vector genome copy numbers are expressed as double-stranded (ds) vector genome copy numbers per diploid genomic equivalent (ds-vg/dge). Error bars represent SEM. A two-way ANOVA followed by Tukey’s post hoc test was used for statistical assessment of the data. ****, adjusted p<0.0001; ns, not significant. **(C)** Representative images of native tdTomato fluorescence in the kidney. Scale bar, 1 mm. Please note that, for the RV and RP groups, the data obtained from the AAV vector-treated kidneys, not non-treated kidneys in the same animals, are presented.

**Figure 3.**
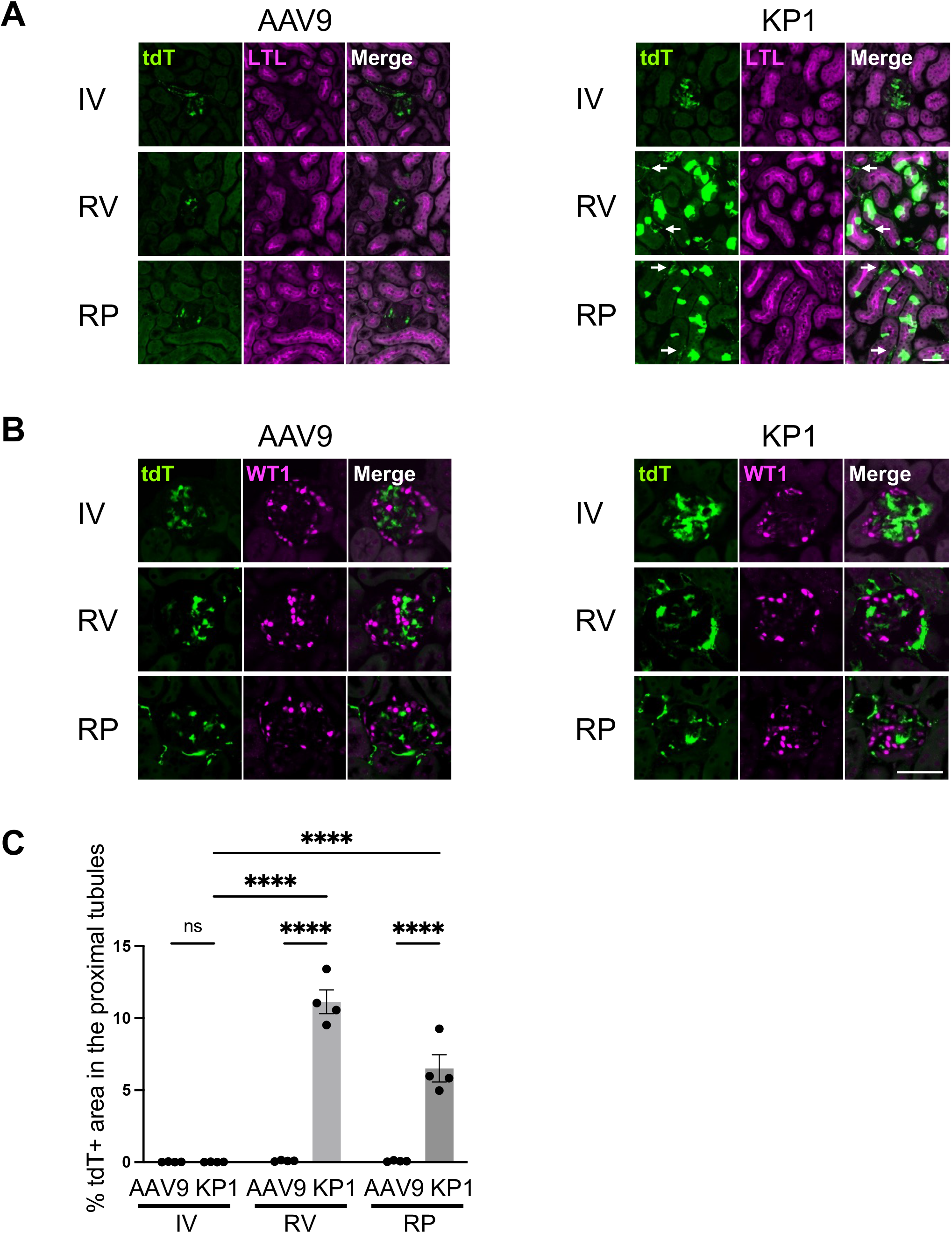
Enhanced transduction was observed in proximal tubule cells by RV or RP injection of AAV-KP1, but not observed in podocytes. Eight-week-old C57BL/6J male mice were injected with AAV9-CAG-tdTomato or AAV-KP1-CAG-tdTomato via RV or RP at a dose of 3x10^11^ vg/mouse (n=4 per group). Kidneys were harvested 2 weeks post injection for downstream analyses. **(A, B)** Representative fluorescence microscopic images of the AAV vector-treated mouse kidneys. In Parel A, green and magenta colors represent native tdTomato fluorescence and LTL (a marker for proximal tubules), respectively. In Panel B, green and magenta colors represent native tdTomato fluorescence and WT1 (a marker for podocytes), respectively. White arrows indicate transduction of interstitial cells. LTL, tetragonolobus lectin; WT1, Wilms tumor 1. Scale bars, 50 μm. **(C)** AAV9 and AAV-KP1 transduction efficiencies in the proximal tubules. The transduction efficiencies were assessed by quantifying tdTomato positive areas in the proximal tubules labeled with LTL. Fiji version of ImageJ was used for the image analysis. A two-way ANOVA followed by Tukey’s post hoc test was used for statistical assessment of the data. ****, adjusted p<0.0001; ns, not significant. Error bars represent SEM.

### Ischemia does not enhance renal transduction of AAV-KP1

Since we performed RV and RP injections with the 15-min blockage of the local arterial and venous flow, we investigated if ischemia influences renal transduction. IV injection of AAV-KP1-CAG-tdTomato (3x10^11^ vg/mouse) into eight-week-old C57BL/6J male mice following 15-min renal ischemia did not alter the transduction, showing only mesangial cell transduction (**Figure S2A**). This indicates that ischemia is not the cause of enhanced renal transduction observed using either RV or RP injections. We also performed RV and RP injections of AAV-KP1-CAG-tdTomato (3x10^11^ vg/mouse) with varying ischemic time periods from 0 min up to 15 min (**Figure S2B)**. There was no apparent difference in transduction efficiency by RV between the following four ischemic time periods (<1 min, 5 min, 10 min and 15 min). Please note that testing RV in no ischemic time period condition is not possible because the renal blood flow needs to be stopped when the agent is injected via RV to the kidney. In the shortest ischemic time period condition for RV injection (*i.e.*, <1 min in **Figure S2B**), we stopped the renal blood flow for no more than 1 min. On the other hand, in the RP injection group, while there was no difference in transduction efficiency between the 5 min, 10 min and 15 min time period groups, RP injection without blockage of renal artery and vein resulted in lower transduction than that in the other time period groups, showing no proximal tubule transduction. Our findings suggest that there is no benefit of prolonged ischemia on kidney cell transduction, but the stoppage of the renal circulation at the time of injection is a crucial step in the RP injection procedure.

### Augmented renal interstitial accumulation of vector particles is a potential mechanism of the enhanced renal transduction by local vector administration

When AAV vectors are administered intravascularly, vascular endothelial cells pose a significant physical and functional barrier for *in vivo* AAV vector transduction in the organs whose endothelium is devoid of fenestrae (*i.e.*, continuous endothelium) such as the brain, heart and skeletal muscles.^19^ In the kidney, endothelial cells of cortical peritubular capillaries and glomeruli are both fenestrated by holes of approximately 60 to 80 nm in diameter with the former having a diaphragm and the latter being devoid of it.^26^ In addition, there is no basement membrane between the glomerular endothelial cells and the mesangial interstitial space in contrast to the peritubular capillary and the other renal blood vessels that have basement membrane.^27, 28^ Based on these microanatomical structure of the kidney, we reasoned that the mesangial interstitial space is readily reached by intravascularly injected AAV vector particles through the fenestrae in the glomerular endothelial cells. The peritubular interstitium in the cortex is also presumed to be reached by intravascularly injected AAV vector particles relatively easily due to the fenestrated nature of the peritubular capillaries, although not as easily as the mesangial interstitial space due to the presence of a diaphragm and basement membrane.

To investigate the mechanism of enhanced transduction by RV and RP injections, we assessed the distribution and extravasation of AAV vector particles in the kidney following AAV vector injection by IV, RV and RP. To this end, we employed fluorescent microspheres of 25 nm in diameter to mimic AAV vector particles. This size of the microspheres was chosen as it is close to the diameter of AAV virions. After 30-min circulation, microspheres administered intravenously mainly accumulated in the mesangial area of the glomeruli (**Figure S3A)**, consistent with the previous findings that nanoparticles accumulate in the mesangial area in a size-dependent manner.^29, 30^ This observation is in line with our knowledge of the above-described anatomical structure of the kidney and the observed effective transduction of mesangial cells with AAV vectors. On the other hand, limited extravasation of microspheres from the peritubular capillary to the interstitial space was observed (indicated by arrows in **Figure S3A)**. Next, we administered microspheres via RV and RP to assess their extravasation and accumulation in the kidney. Interestingly, both administration methods resulted in efficient interstitial accumulation of the microspheres, predominantly in the renal cortex (**Figure S3B and S3C)**. It is important to note that interstitial accumulation was not observed by RP injection in the absence of the blockage of renal artery and vein (**Figure S3D**). These observations suggest that the peritubular capillary wall poses a significant barrier for AAV vector to transduce tubular epithelial cells following IV injection even with the presence of fenestration in the endothelium, while RV and RP injections allow AAV vector particles to pass through or bypass this barrier, leading to efficient transduction of the proximal tubules. The lack of transduction enhancement by RP injection without local blockage of the blood flow (**Figure S2B**) also supports the importance of interstitial accumulation of AAV vector particles for proximal tubule transduction.

### Limited extra-renal dissemination of AAV-KP1 following local injection suppresses off-target liver transduction

Several studies have reported off-target organ transduction, especially in the liver, following local injection of AAV9.^13, 31–33^ Therefore, we next assessed the off-target liver transduction following RV and RP injections of the AAV9 and AAV-KP1 vectors. Quantification of vector genomes in the liver two weeks after IV, RV and RP injections revealed that both AAV9 and AAV-KP1 are equally liver tropic by IV injection, while RV and RP injections of the AAV-KP1 reduced the off-target liver transduction by 14-fold and 7-fold compared to IV injection, respectively (two-way ANOVA followed by Tukey’s post hoc test, adjusted p=5.8x10^-6^ for RV and adjusted p=6.7x10^-4^ for RP). On the other hand, off-target liver transduction was not prevented by RV and RP injections of AAV9 (**Figure 4A**). A fluorescence microscopic analysis also confirmed substantially suppressed off-target liver transduction by RV and RP injections of AAV-KP1 compared to AAV9 (**Figure 4B**). Based on these observations, we hypothesized that the difference in pharmacokinetics following local injection leads to the contrasting renal and hepatic transduction between AAV9 and AAV-KP1. Quantification of vector genomes 10 min after local injection showed >10 times more vector genomes in the AAV-KP1-injected kidney than the AAV9-injected kidney following RV and RP (**Figure 4C**) injections. This finding indicates that injected AAV-KP1 vector particles stay in the kidney more efficiently than AAV9 vector particles. It is important to note that transduction with AAV vector does not occur within 10 min following vector exposure *in vivo.*^19^ Thus, it is unlikely that the higher concentration of AAV-KP1 vector genomes than AAV9 vector genomes in the injected kidney is the consequence of higher transduction of the AAV-KP1-injected kidney. Rather, we postulate that AAV-KP1 vector particles were trapped by the renal interstitium more efficiently than AAV9 vector particles following injection, and this accounts for the observed difference in AAV9 and AAV-KP1 vector genome copy numbers in the kidney in the very early stage of *in vivo* transduction. In addition, blood concentrations of AAV-KP1 were more than 50 times less than those of AAV9 throughout the study time points (0 min, 10 min, 30 min, 1 h, 4 h and 8 h after RV and RP injections, **Figure 4D**). These observations have led us to a model that explains the mechanism for the significant enhancement of on-target kidney transduction and suppression of off-target organ transductions by local injection of AAV-KP1, and the failure of AAV9 to achieve similar outcomes. In this mechanistic model, the unique biological attribute of AAV-KP1 that allows its effective sequestration within the kidney enables a significant increase in the local concentration of AAV vector particles, thereby substantially enhancing renal transduction. In the meantime, this attribute limits the spillover of vector particles from the kidney, minimizing vector dissemination to off-target organs. On the other hand, AAV9, lacking this attribute, rapidly exits the kidney without accumulating upon local injection and effectively enters the bloodstream, resulting in significant vector spillover in off-target organs. Therefore, AAV9 is not an appropriate AAV capsid for local vector delivery with limited off-target effects.

**Figure 4.**
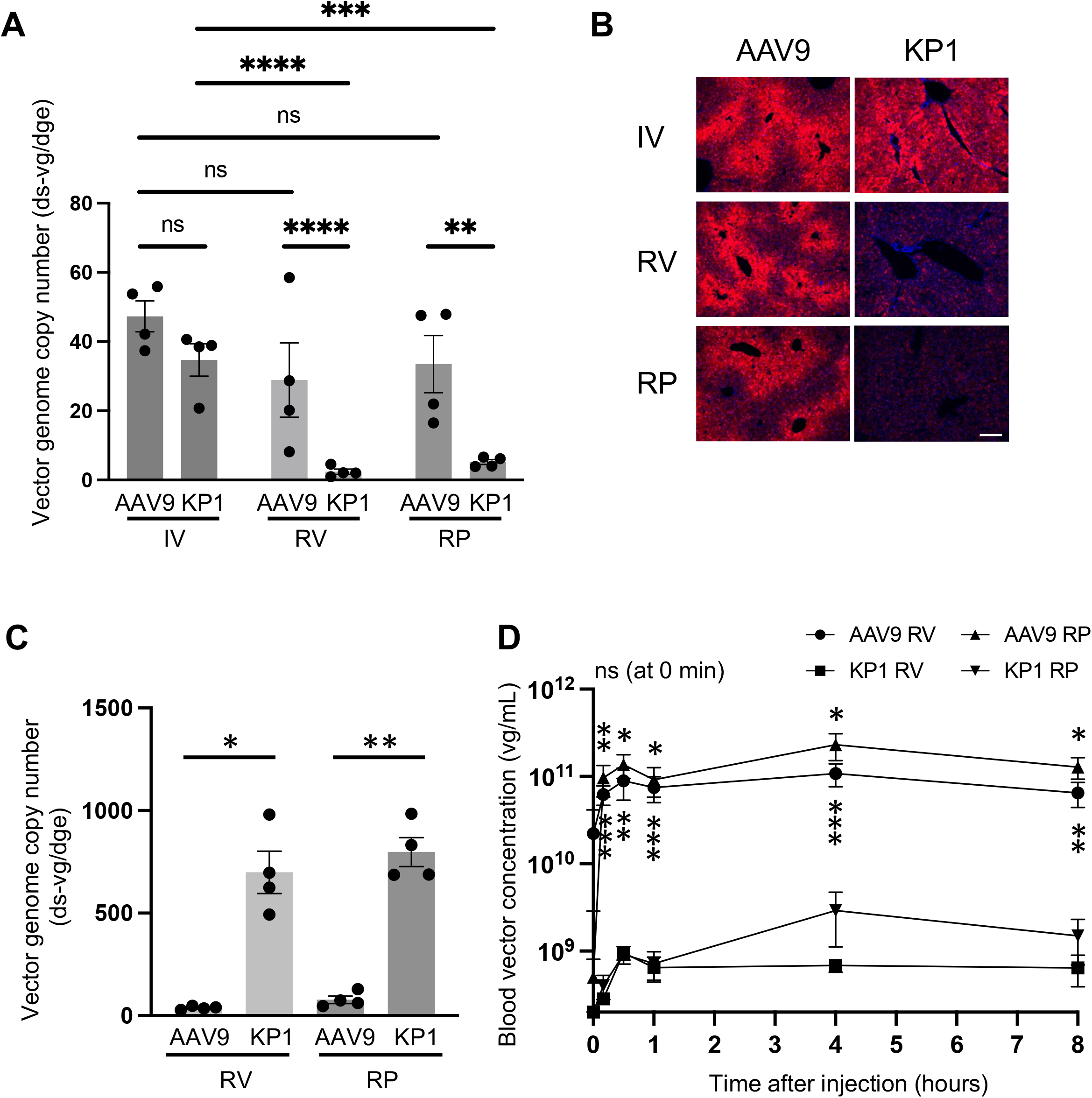
AAV vector pharmacokinetics following RV and RP injections and their implications for off target liver transduction. Eight-week-old C57BL/6J male mice were injected with AAV9-CAG-tdTomato or AAV-KP1-CAG-tdTomato via IV, RV or RP at a dose of 3x10^11^ vg/mouse (A, B) or 1x10^13^ vg/kg (C, D) (n=4 per group). **(A)** AAV vector genome copy numbers in the livers harvested 2 weeks post-injection were determined by qPCR. Vector genome copy numbers are expressed as double-stranded (ds) vector genome copy numbers per diploid genomic equivalent (ds-vg/dge). **(B)** Representative fluorescence microscopic images of the livers in the AAV vector-treated mice. Red, native tdTomato fluorescence; blue, DAPI. Scale bar, 100 μm. **(C)** AAV vector genome copy numbers in the kidneys in the AAV vector-treated mice were determined by qPCR 10 min after RV and RP injections. **(D)** Blood vector concentration-time curves following RV and RP injection. A two-way ANOVA followed by Tukey’s post hoc test was used for statistical assessment of the data presented in Panel A. In Panel C and D, a two-tailed Welch’s t-test with Bonferroni correction was used. In Panel D, the p value represents the statistical significance between AAV9 and AAV-KP1 by RV or RP injection at each time point. Please note that there was no significant difference between AAV9 and AAV-KP1 at 0min by both injection methods. ****, adjusted p<0.0001; ***, adjusted p<0.001; **, adjusted p<0.01; *, adjusted p<0.05; ns, not significant. Error bars represents SEM.

### Urinary excretion of AAV9 is substantially increased in CKD

To successfully apply AAV vector approaches for the treatment of CKD, it is crucial to understand AAV vector transduction and pharmacokinetic profiles in CKD. To this end, we employed the Col4a5 mutant mouse model harboring a G5X nonsense mutation in the α5 chain of type IV collagen (B6.Cg-Col4a5^tm1Yseg^/J),^34^ which serves as a valuable mouse model of X-linked Alport syndrome. Hemizygous male mice of this strain manifest progressive renal failure due to Alport syndrome, which is the second common genetic cause of CKD in adults.^6^ To characterize *in vivo* AAV vector biology in this CKD mouse model, we injected 25 to 30-week-old B6.Cg-Col4a5^tm1Yseg^/J hemizygous male mice and age-matched wild-type mice from the colony with AAV9-CAG-tdTomato or AAV-KP1-CAG-tdTomato vector at a dose of 1x10^13^ vg/kg (approximately 3x10^11^ vg/mouse, n=3 to 5 per group, **Figure 5A**). As reported previously,^34^ our hemizygous mutant male mice manifested significant albuminuria. Urinary albumin excretion of the mutant mice and the wild-type controls at the time of injection was 1.8 ± 0.14 g/gCre and 0.047 ± 0.0098 g/gCre, respectively (two-tailed Welch’s t-test, p=3.4x10^-7^, **Figure 5B**), suggesting increased permeability of the glomerular filtration barrier. To investigate whether the altered glomerular filtration barrier affects the pharmacokinetics of AAV vectors, we quantified urinary excretion of AAV vector particles over 5 hours following IV injection of the AAV9 or AAV-KP1 vector. In the wild-type control mice, only about 0.001% of injected AAV9 and AAV-KP1 vector particles defined by vector genome titers were excreted into the urine during the 5-hour period (4.7x10^5^ to 1.5x10^6^ vg of a total of 3 x 10^11^ vg injected into each animal, **Figure 5C**). In contrast, more than 1% of injected AAV9 vector particles (3.8x10^9^ to 9.5x10^9^ vg of a total of 3 x 10^11^ vg injected into each animal) were excreted, showing a 10^4^ times increase compared to the wild-type control mice (two-way ANOVA followed by Tukey’s post hoc test, adjusted p=2.7x10^-5^, **Figure 5C**). As it is known that leaky glomerular filtration barrier and decreased renal tubular reabsorption contribute to albuminuria in CKD, our results suggest that the glomerular filtration barrier and potentially renal tubular reabsorption play an important role in preventing the urinary excretion of AAV vectors. Interestingly, unlike AAV9, the urinary excretion of AAV-KP1 in the CKD mouse model was only modestly increased with no statistical significance (two-way ANOVA followed by Tukey’s post hoc test, adjusted p=0.10, **Figure 5C**). Although glomerular filtration is not solely determined by blood concentration, blood and urine concentrations of substances are positively correlated in many instances;^35^ therefore, it is reasonable to hypothesize that the contrasting urinary excretion profiles of AAV9 and AAV-KP1 are primarily attributed to the difference in their blood concentrations. To support this, AAV9 and AAV-KP1 showed distinct pharmacokinetics over 8 hours following IV administration. In contrast to the delayed blood clearance of AAV9, which we previously reported,^19^ AAV-KP1 was rapidly cleared from the bloodstream following IV administration, resulting in 48 times less blood concentration at 1 min and 572 to 2152 times less blood concentrations at the time points between 30 min and 8 h post-injection (**Figure S4**). These results provide indirect evidence that urinary excretion of AAV vector particles is mainly prevented by the glomerular filtration barrier rather than tubular reabsorption in the healthy kidney.

**Figure 5.**
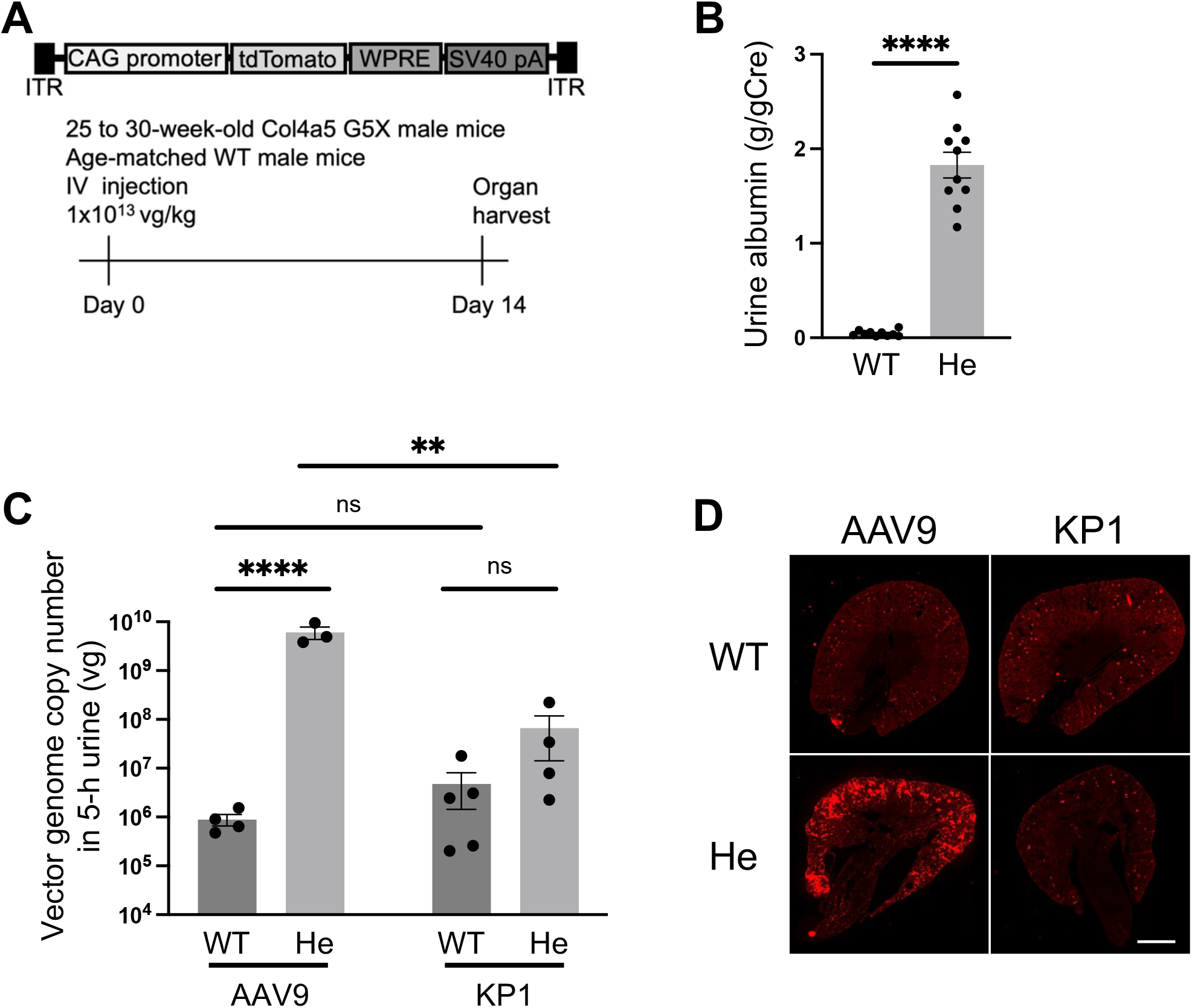
Enhanced AAV-vector mediated renal transduction and urinary AAV excretion in CKD following IV injection of AAV9. **(A)** The experimental design. Twenty-five to 30-week-old Col4a5 G5X hemizygous male mice (He) and age-matched wild-type control male mice (WT) were injected with AAV9-CAG-tdTomato and AAV-KP1-CAG-tdTomato via IV at a dose of 1x10^13^ vg/kg. Kidneys were harvested 2 weeks post-injection for downstream analyses. **(B)** Albumin levels in the urine samples collected from the 25 to 30-week-old He and WT mice (n=10 each) at the time of injection. The data was normalized by urine creatinine concentration and expressed as g/gCre. A Welch’s t-test was used for statistical assessment. **(C)** Urinary excretion of AAV-CAG-tdTomato vectors over 5 hours following IV injection. AAV vector genome copy numbers in urine samples were determined by qPCR in the following 4 groups. AAV9 in WT (n=4), AAV9 in He (n=3), AAV-KP1 in WT (n=5) and AAV-KP1 in He (n=4). A two-way ANOVA followed by Tukey’s post hoc test was used for statistical assessment of the data. ****, p<0.0001 or adjusted p<0.0001; **, adjusted p<0.01; ns, not significant. Error bars represent SEM. **(D)** Representative fluorescence microscopic images of the kidneys in the AAV-vector treated mice. Red, native tdTomato fluorescence; blue, DAPI. Scale bar, 100 μm.

### IV injection of AAV9 transduces podocytes and proximal tubule cells effectively in the CKD kidney

We next analyzed the renal transduction efficiency and cell type tropism of AAV9 and AAV-KP1 in the CKD mouse model. Consistent with the urinary excretion data, global renal transduction of the AAV9 vector was substantially enhanced in the CKD kidney, which was not observed for in the CKD mice injected with the AAV-KP1 vector (**Figure 5D**). The enhanced transduction of AAV9 in the CKD kidney was mainly observed in the cortex. We then conducted co-staining analyses using cell type-specific markers to determine the identity of the transduced cell types. Co-staining of proximal tubule cells revealed that AAV9 transduced 9.4% of proximal tubule cells of the CKD kidney while no proximal tubule transduction was observed in the wild-type control kidney, demonstrating remarkably enhanced proximal tubule transduction with AAV9 in the CKD kidney (**Figure 6A, 6C**). On the other hand, proximal tubule transduction was not observed in the CKD mice injected with the AAV-KP1 vector (**Figure 6A, 6C**). Moreover, co-staining of podocyte nuclei revealed that AAV9 and AAV-KP1 transduced 35% and 15% of podocytes in the CKD kidney (**Figure 6B, 6D**). The remarkable improvement of podocyte transduction in the CKD kidney following IV administration of AAV vectors aligns with our hypothesis that the increased permeability of the glomerular filtration barrier in the CKD kidney renders podocytes more accessible to AAV vectors following IV delivery. The higher podocyte transduction with AAV9 than AAV-KP1 (two-way ANOVA followed by Tukey’s post hoc test, adjusted p=0.011) is consistent with the higher urinary excretion of AAV9 than AAV-KP1 in the CKD mice. We also investigated cardiac transduction with AAV9 and AAV-KP1. We hypothesized that the cardiac transduction with AAV vectors could be affected in the CKD mice as CKD has a systemic effect especially on cardiovascular organs^36^ and cardiac remodeling such as hypertrophy, fibrosis, capillary rarefaction is a common feature of CKD including Alport syndrome.^37–39^ The results showed significantly enhanced cardiac transduction with AAV9 in the CKD mice compared to the wild-type mouse controls (two-way ANOVA followed by Tukey’s post hoc test, adjusted p=0.018, **Figure S5**). On the other hand, there was no significant difference in cardiac transduction between the CKD mice and the wild-type controls injected with AAV-KP1 (two-way ANOVA followed by Tukey’s post hoc test, adjusted p=0.99, **Figure S5**). AAV-KP1 exhibited lower cardiac tropism than AAV9 (two-way ANOVA followed by Tukey’s post hoc test, adjusted p=0.046, **Figure S5**). Therefore, no alteration in cardiac transduction with AAV-KP1 in the CKD mice could be attributed to the inherent absence of cardiac tropism in AAV-KP1. These observations underscore the importance of the host condition as a key determinant of AAV vector transduction, with varying effects among different AAV capsids.

**Figure 6.**
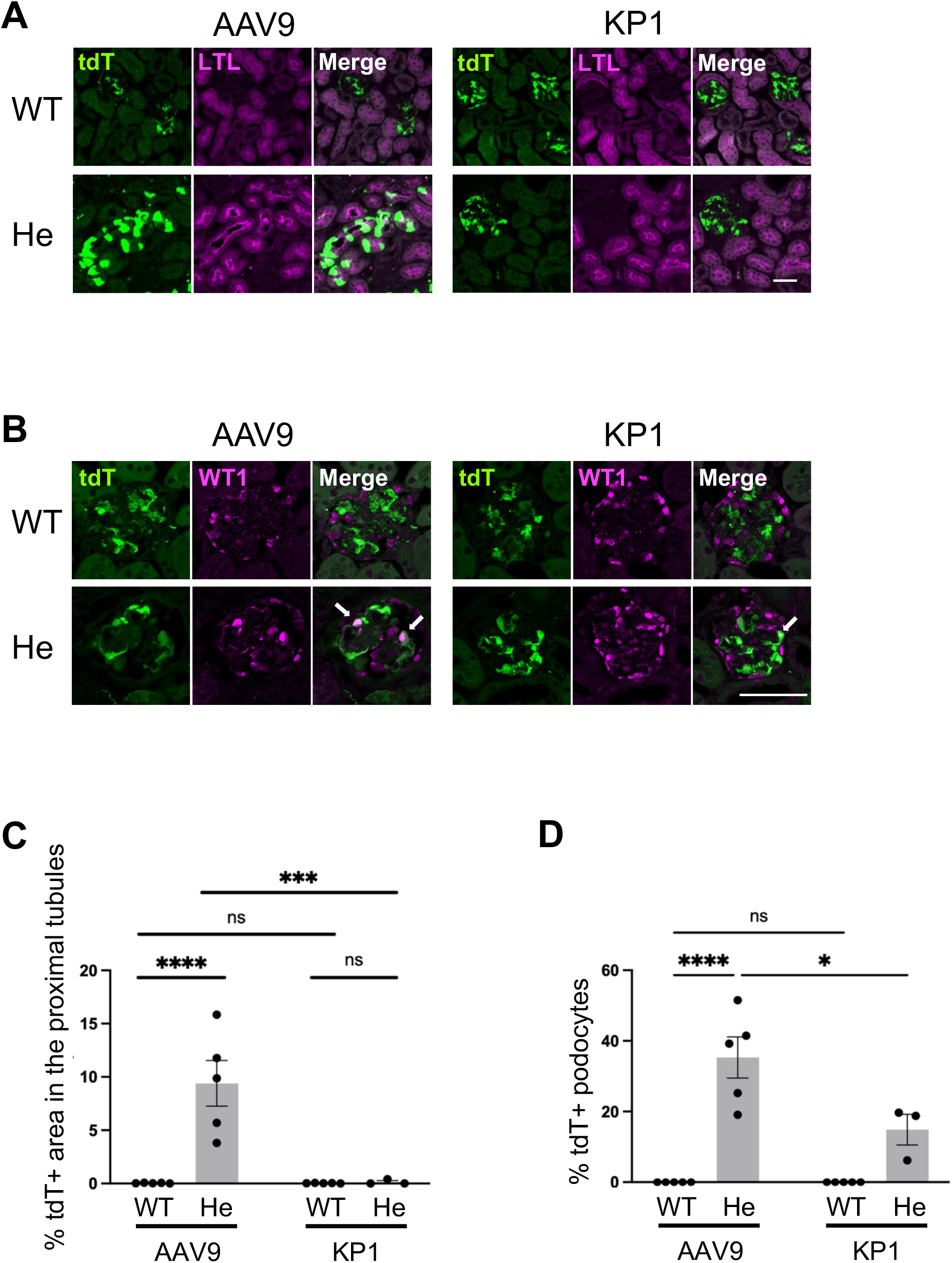
Enhanced AAV9-mediated transduction of proximal tubule cells and podocytes in CKD following IV injection. Twenty-five to 30-week-old Col4a5 G5X hemizygous male mice (He) and age-matched wild-type control male mice (WT) were injected with AAV9-CAG-tdTomato and AAV-KP1-CAG-tdTomato via IV at a dose of 1x10^13^ vg/kg. Kidneys were harvested 2 weeks post-injection for downstream analyses. AAV9 in WT (n=5), AAV9 in He (n=5), AAV-KP1 in WT (n=5) and AAV-KP1 in He (n=3). **(A, B)** Representative fluorescence microscopic images of the kidneys in the AAV vector-treated mice. In Parel A, green and magenta colors represent native tdTomato fluorescence and LTL (a marker for proximal tubules), respectively. In Panel B, green and magenta colors represent native tdTomato fluorescence and WT1 (a marker for podocytes), respectively. White arrows indicate AAV vector-transduced podocytes. Scale bars, 50 μm. **(C)** AAV9 and AAV-KP1 transduction efficiencies in the proximal tubules. The transduction efficiencies were assessed by quantifying tdTomato positive/negative areas in the proximal tubules labeled with LTL. Fiji version of ImageJ was used for the image analysis. **(D)** AAV9 and AAV-KP1 transduction efficiencies in podocytes. The transduction efficiencies were assessed by a manual counting of tdTomato-positive/negative podocytes labeled with WT1. A two-way ANOVA followed by Tukey’s post hoc test was used for statistical assessment of the data. ****, adjusted p<0.0001; ***, adjusted p<0.001; *, adjusted p<0.05; ns, not significant. LTL, tetragonolobus lectin; WT1, Wilms tumor 1.

## Discussion

In this study, we have identified a small subset of AAV capsids that exhibit significantly enhanced gene transfer to the kidney by local vector administration. By using the AAV capsids showing this peculiar phenotype, we could transduce proximal tubules at levels that has not been attainable before. According to the previous studies assessing renal transduction of common AAV serotypes by IV,^10, 11^ RV^12^ and RP^14–16, 40, 41^ injections, it has been generally considered that AAV9 is one of the most efficient capsids regardless of administration routes. However, our unbiased, head-to-head comparison study that sought to determine renal transduction efficiency with various AAV capsids covering both common serotypes and engineered capsids has demonstrated that effective AAV capsids are not necessarily the same between systemic IV injection and local injections. We show here that remarkable enhancement of renal transduction can be attained by local injection only when a particular subset of AAV capsids is used. In the study, we have identified the following six AAV capsids, AAV-KP1, AAV-KP2, AAV-KP3, AAV-DJ, AAV2G9 and AAV2.7m8, as those that outperform AAV9 only when administered by RV or RP injection. AAV9 is the serotype that shows the most robust *in vivo* transduction in various organs following IV injection and our AAV Barcode-Seq experiment has also confirmed it in the context of renal gene transfer. AAV-KP and AAV-DJ^22^ were generated by capsid shuffling and *in vitro* screen in pancreatic islet and hepatoma cells, respectively. AAV2G9 was generated by engraftment of the galactose binding motif in the AAV9 capsid into the AAV2 capsid.^23^ AAV2.7m8 was generated by an *in vivo* screen of AAV capsid mutants in photoreceptors of the mouse retina.^24^ Therefore, their high performance in renal gene transfer by local administration was unexpected and not predictable, exemplifying the utility of high-throughput AAV capsid phenotype characterization such as AAV Barcode-Seq. Our discovery that there is a small subset of AAV capsids that can substantially enhance renal transduction by local vector administration represents a breakthrough in renal gene transfer and opens a new avenue to the development of much more effective approaches for gene therapy to treat kidney diseases.

One of the most important findings in our study is that AAV-KP1 efficiently transduces the proximal tubule cells by local vector administration. The proximal tubules account for a half of the tubular epithelial cells^42^ and is an important target cell type for both basic research^43^ and gene therapy.^8^ Therefore, the ability of AAV-KP1 to target proximal tubule is a promising feature. It is important to note that RV and RP injections are not parenchymal injections that could damage tissues. It is also important to emphasize that RV or RP-injected AAV vectors can infiltrate into the whole kidney, resulting in widely distributed proximal tubule transduction with AAV vectors. Furthermore, we showed that the extended blockage of the renal arterial and venous flow performed in previous studies^12, 15, 16^ is not required for efficient renal transduction of AAV-KP1. Given the difficulties in achieving widespread transduction with AAV vectors by local vector delivery in other organs such as the brain, our observation underscores the kidney as a compelling target organ for local gene delivery approaches to effectively treat diseases. In addition, the ability for RV and RP injections to achieve effective renal gene transfer while restricting the duration of blood flow blockage solely to the time required for completing the injections holds significant promise for clinical translation of the local approaches for renal gene therapy.

To further enhance the attractiveness and clinical relevance of the local renal gene delivery approaches, we have discovered AAV-KP1 as an AAV capsid exhibiting a compelling phenotype that mitigates off-target effects. This capsid demonstrates a unique phenotype that limits vector dissemination to off-target organs and achieves highly efficient and specific transduction of renal cells. Recent clinical trials raised concerns about adverse effects related to the dissemination of AAV vectors to off-target organs, including hepatotoxicity, genotoxicity, thrombotic microangiopathy and dorsal root ganglia (DRG) toxicity.^1, 17^ Local administration of AAV-KP1 has a potential to prevent these side effects of AAV vector-mediated gene therapy. We demonstrated in this study that the degree of vector spillover following local administration is capsid dependent and varies significantly among different AAV capsids. The local administration of AAV9 into the kidney resulted in liver transduction at levels comparable to those achieved by systemic administration. This finding indicates that AAV9 is not suitable for local injection when an approach also aims to minimize off-target effects. Importantly, these phenotypes of AAV9 and AAV-KP1 are not specific to the kidney. For example, we observed contrasting patterns of target and off-target organ transductions between AAV9 and AAV-KP1 following pancreatic duct injection in rhesus macaques. In this non-human primate study, AAV-KP1 also showed high pancreatic transduction while minimizing off-target liver transduction (Adachi *et al.* American Society of Gene and Cell Therapy 25^th^ annual meeting, Washington D.C., 2022). Substantial off-target liver transduction was also previously reported by other groups following local administration of AAV9 to the cerebrospinal fluid^31, 32^ and pancreatic duct.^33^ These observations suggest that AAV-KP1 has an inherent phenotype that allows for accumulation at the injection site, thereby enabling efficient renal transduction and minimizing vector spillover to off-target organs. Studies in other organs will be needed to further characterize this unique phenotype of AAV-KP1.

Based on all of our observations including the microsphere distribution in the kidney, we propose a model in which the microanatomical structures of capillary endothelia and basement membranes in the kidney and the interstitial accumulation of AAV vector particles play pivotal roles in mediating efficient transduction in mesangial cells following IV injection and efficient transduction of the proximal tubules following RV and RP injections (**Figure 7**). This model also explains the challenges in AAV vector transduction in the kidney. As mentioned in the RESULT section, the fenestrae of the glomerular endothelial cells lack a diaphragm,^26^ and there is no basement membrane between the glomerular capillary and the mesangium.^27, 28^ This unique architecture facilitates efficient mesangial transduction following IV injection. On the other hand, the pathway from the peritubular capillary to the tubular epithelial cells is composed, in series, of a fenestrated endothelium with a diaphragm, the capillary basement membrane, the interstitium and the tubular basement membrane.^29, 44^

**Figure 7.**
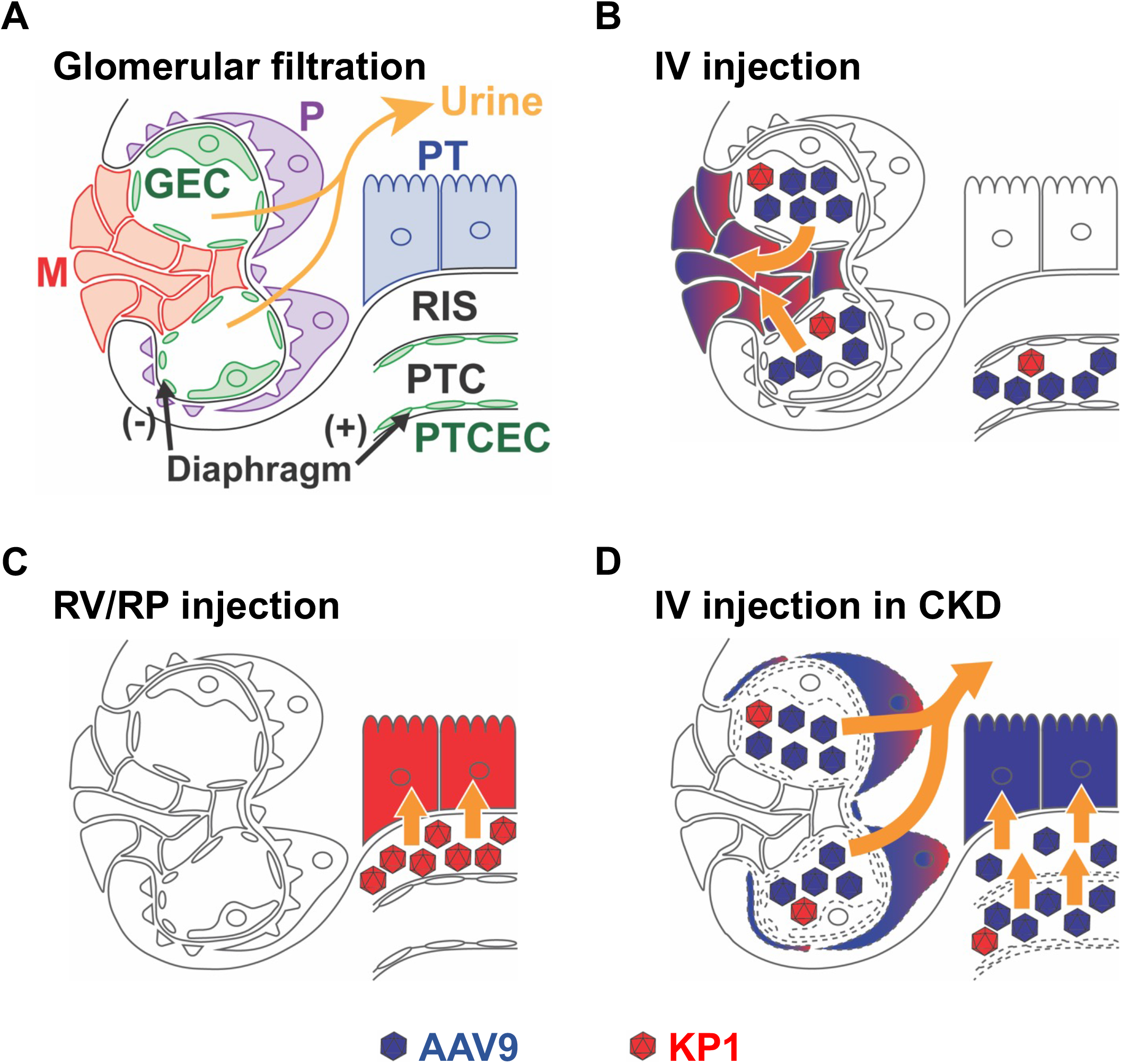
A schematic summary illustrating AAV9 and AAV-KP1-mediated renal transduction by IV, RV and RP injections in healthy and CKD conditions. **(A)** Glomerular filtration occurs across the glomerular endothelial cells (GEC), the glomerular basement membrane and podocytes (P). The filtration flow is indicated with yellow arrows. The renal interstitial space (RIS) is the space between the proximal tubules (PT) and the peritubular capillaries (PTC). M, mesangial cell; PTCEC, peritubular capillary endothelial cell. Please note that glomerular filtration of AAV is prevented in the healthy kidney by the glomerular filtration barrier. **(B, C and D)** A mechanistic model illustrating the transduction of proximal tubules and podocytes with AAV9 (blue) and AAV-KP1 (red) by IV, RV and RP injections. Cells transduced with AAV9 and AAV-KP1 are highlighted in colors corresponding to their respective AAV capsids. **(B)** Upon IV injection, AAV is preferentially distributed into the mesangial area (M) through the fenestrae of GEC that lacks both a diaphragm and a basement membrane. Note that GEC has a basement membrane except for the GEC in the mesangial area. **(C)** By RV and RP injections, AAV can reach RIS. However, only AAV-KP1, not AAV9, can accumulate in RIS due to its unique attribute and achieves proximal tubule transduction. **(D)** Upon IV injection in CKD, the impaired GEC and PTCEC (indicated by dashed lines) enhance AAV excretion into the urine and promote AAV extravasation into RIS due to the increased permeability of the glomerular filtration barrier and the PTCEC barrier, respectively. This enables enhanced transduction of podocytes and proximal tubules cells following IV injection. Both AAV9 and AAV-KP1 transduce podocytes, with AAV9 exhibiting higher transduction efficiency due to its increased concentration in the urine (Figure 5C). On the other hand, only AAV9, not AAV-KP1, transduces proximal tubules because, unlike AAV9, AAV-KP1 fails to accumulate in RIS most likely due to its rapid blood clearance following IV injection. Mesangial cell transduction occurs following both RV and RP injections and IV injection in CKD, but this is not depicted in this schematic.

This architecture makes tubular epithelial cell transduction by IV more challenging. In the RV and RP injections, both AAV9 and AAV-KP1 are distributed to the interstitial space; however, only AAV-KP1 remains within the interstitial space efficiently and transduces the proximal tubule cells from the basolateral surface without undergoing rapid redistribution to the systemic circulation via venules or the lymphatic system. The idea that RV and RP injections facilitate the proximal tubule transduction from the basolateral interstitial side is supported by our AAV Barcode-Seq data. Although RV and RP administration routes are anatomically distinct, our AAV Barcode-Seq analysis showed similar transduction profiles across all the 47 different AAV capsids and identified the same six AAV capsids that show remarkably enhanced renal transduction by RV and RP injections, suggesting that the transduction pathway is shared by these two injection methods. The shared pathway in AAV vector-mediated renal tubule transduction by RV and RP is likely the interstitial accumulation and the subsequent transduction of renal tubule cells from the basolateral side. To support this, RV and RP injections of AAV-KP1 also resulted in interstitial transduction (indicated by arrows in **Figure 3A**). Please note that our model does not address the mechanism by which AAV vector particles accumulate in the renal interstitium following RV and RP injections. In RV injection, the mechanistic understanding could be straightforward because the increased hydrostatic pressure in the peritubular capillary lumen likely facilitates the extravasation of AAV vector particles, resulting in the accumulation of AAV vector particles in the interstitium. On the other hand, the mechanism of the interstitial accumulation of AAV vector particles following RP injection is less straightforward. Presumably, AAV vector particles enter the interstitium from the urinary lumen via paracellular pathways that are transiently opened by the pressure generated during the RP procedure. Further studies will be needed to better understand these mechanisms.

Our study demonstrates that AAV vector-mediated efficient transduction of podocytes and proximal tubules at levels of 35% and 9.4%, respectively, can be achieved in the CKD mouse model by IV injection of AAV9 at a dose of 1 x 10^13^ vg/kg. A previous study used AAV9 in a Col4a3 deficient mice, a mouse model of autosomal recessive Alport syndrome. However, this study did not assess differences in transduction between healthy and diseased kidneys; therefore, the potential effects of CKD on AAV transduction remained unknown.^45^ Interestingly, AAV9 outperforms AAV-KP1 in the transduction of both cell types in the CKD kidney following IV injection. The demonstration of the ability of AAV9 to transduce approximately one-third of podocytes holds immense clinical implications, given that podocytes represent the most common target cell type for monogenic kidney diseases^8, 46^ and AAV9 has been used as a clinical IV product approved by regulatory agencies. Although detailed mechanisms for enhanced podocyte transduction in CKD will be the focus of future studies, our data suggest that the increased permeability of the glomerular filtration barrier plays a crucial role. The difference in proximal tubule transductions in CKD between AAV9 and AAV-KP1 is presumably explained by the difference in their pharmacokinetic profiles. A previous study has shown that the peritubular capillary undergoes functional and structural changes in CKD, with the increased permeability of the peritubular capillary being a common feature of CKD, including Alport syndrome.^18^ Thus, in the CKD kidney, AAV9’s delayed blood clearance should provide a greater opportunity for a larger quantity of vector particles to pass through the compromised peritubular capillary wall compared to AAV-KP1. This disparity in the pharmacokinetics presumably allows AAV9 to transduce more proximal tubule cells than AAV-KP1 from the interstitial basolateral side **(Figure 7**). Although proximal tubule transduction from the interstitial basolateral side is consistent with the proposed mechanistic model in healthy kidney described earlier, it is also possible that AAV9 with increased concentration in the urinary lumen in CKD mice transduces the proximal tubules from the apical side. High concentration of AAV9 vector particles in the urinary lumen delivered by RP injection in healthy kidney did not result in proximal tubule transduction, but proximal tubule cells in CKD may undergo phenotypic changes that render the cells susceptible to transduction with AAV9.

Our data showed that CKD altered the AAV vector transduction profiles in the kidney and remote organs such as the heart. Our knowledge about potential health condition-dependent variations in AAV transduction profiles remains limited. In acute diseases such as epilepsy^47^ and myocardial infarction,^48^ enhanced AAV transduction has previously been reported. However, there is a paucity of knowledge about potential changes in AAV vector transduction profiles in the context of chronic diseases, which represent the most common health conditions in AAV vector-mediated gene therapy. Our CKD study conveys two important messages. First, in the context of AAV vector-mediated renal gene therapy, it is crucial to acknowledge that disease-associated pathophysiological changes can influence AAV vector transduction profiles. Consequently, a meticulous selection of both AAV capsids and the route of administration, tailored to the health condition of the kidney, becomes paramount. Second, higher urinary excretion of AAV vectors and a potential increase in off-target effects in patients with CKD should be fully addressed prior to the clinical implementation of AAV vector-mediated gene therapies for CKD. Because current clinical AAV vector-mediated gene therapies exclude patients with CKD,^49–51^ our clinical experience regarding the use of AAV in CKD is currently lacking.

In summary, the present study highlights the importance of AAV capsid selection depending on the route of administration and the host condition in the context of AAV vector-mediated renal gene delivery. Our observations suggest that different pharmacokinetics among AAV capsids play an important role in context-dependent performance of AAV vectors. Further research on the context-dependent capsid engineering and characterization is imperative for the development of more effective and safer AAV vector-mediated gene therapy.

## Materials and Methods

### Animal experiments

All the animal experiments were performed according to the guideline for animal care at Oregon Health & Science University (OHSU). C57BL/6J mice and B6.Cg-Col4a5^tm1Yseg^/J mice were purchased from the Jackson Laboratory (Strain IDs are 664 and 6183, respectively). For the AAV Barcode-Seq analysis, eight-week-old C57BL/6J male mice were injected with an AAV barcode library via the tail vein at a dose of 2x10^13^ vg/kg (n=3), and via renal vein (n=4) and renal pelvis (n=3) at a dose of 3x10^11^ vg/mouse. Six weeks post-injection, mice were euthanized, and kidneys were harvested. For the AAV9 and AAV-KP1 individual capsid validation study, eight-week-old C57BL/6J male mice were injected with AAV9-CAG-tdTomato or AAV-KP1-CAG-tdTomato via the tail vein, renal vein, and renal pelvis at a dose of 3x10^11^ vg/mouse (n=4 per group). Two weeks post-injection, mice were euthanized, and kidneys and livers were harvested for vector genome quantification and histological assessment by immunofluorescence microscopy. For a pharmacokinetic study, eight-week-old C57BL/6J male mice were injected with AAV9-CAG-tdTomato or AAV-KP1-CAG-tdTomato via IV, RV or RP at a dose of 1x10^13^ vg/kg (n=4 per group). Subsequently, whole blood samples were collected from the retro-orbital plexus at 6 time points (0 min for RV and RP when 15-min dwelling was completed or 1 min for IV, followed by 10 min, 30 min, 1 h, 4 h and 8 h) following vector injection and vector genome copy numbers in the blood samples were determined. To quantify viral genome copy numbers in the kidney after local vector injection, eight-week-old C57BL/6J male mice were injected with AAV9-CAG-tdTomato or AAV-KP1-CAG-tdTomato via RV or RP at a dose of 1x10^13^ vg/kg (n=4 per group). Subsequently, injected kidneys were harvested 10 min after the completion of 15-min dwelling time following the RV and RP injections and vector genome copy numbers in the kidney were determined. To investigate the renal transduction in CKD, 25 to 30-week-old B6.Cg-Col4a5^tm1Yseg^/J hemizygous male mice and age-matched wild-type controls from the colony were randomly allocated to two groups and intravenously injected with AAV9-CAG-tdTomato and AAV-KP1-CAG-tdTomato at a dose of 1x10^13^ vg/kg (n=3-5 per group). After injection, mice were put in the metabolic cage (MMC100, Hatteras Instruments, Grantsboro, NC) for 5 h to collect the urine samples. Two weeks post-injection, mice were euthanized, and the kidneys and hearts were harvested for downstream analysis.

### Mouse surgical procedures

For renal vein (RV) injection (modified from the previous protocol^12^), mice were anesthetized by isoflurane inhalation and placed on a heated surgical pad (8002062012, Stryker Medical, Portage, MI) to maintain a constant body temperature. A medial abdominal incision was made and intestines were removed from the abdominal cavity to expose the left renal vasculature and the kidney. Removed intestines were kept moist throughout the surgery. A non-traumatic micro-serrefine clamp was placed on the renal artery and vein and 50 ^μ^L of AAV vector solution was injected using a 31-gauge needle (328468, BD Medical, Franklin Lakes, NJ). Ischemic time required for the injection is less than 1 min. Fifteen min post-injection, the clamp was removed to observe the restoration of the blood flow (verified by color change), and the incision was closed. For renal pelvis (RP) injection (modified from the previous protocol^52^), a flank incision was made to expose the left kidney to place a non-traumatic micro-serrefine clamp on the renal pedicle and the ureter under general anesthesia. Fifty μL of AAV solution was injected into the pelvic cavity over 1 min using a syringe pump (70-4507, Harvard Apparatus, Holliston, MA) to prevent parenchymal damage and leakage. Fourteen min post-injection (total ischemic time, 15 min), the clamp was removed to observe the restoration of the blood flow, and the incision was closed. To assess the effect of ischemia on renal transduction, a clamp was placed on the renal artery and vein for 15 min by making a flank incision, and tail vein injection was performed after removal of the clamp. RV and RP injections were also performed with different lengths of ischemia (<1 min (RV injection) or 0 min (RP injection), 5 min, 10 min and 15 min). In the <1 min group, RV injection was performed with a minimal length of ischemia required for the injection. RP injection was performed without the blockage of the renal artery and vein for the 0 min group and a clamp was placed on the ureter for 15 min regardless of the ischemic time.

### AAV vectors and plasmids

AAV-CAG-BC library, AAV9-CAG-tdTomato and AAV-KP1-CAG-tdTomato were produced in HEK293 cells (RRID: CVCL-6871, Agilent) by an adenovirus-free plasmid transfection method and purified by two rounds of cesium chloride density-gradient ultracentrifugation followed by dialysis as described previously.^53^ AAV9 and AAV-KP1 helper plasmids were provided by J. M. Wilson and M. A. Kay, respectively. pAAV-CAG-tdTomato was a gift from E. Boyden (59462, Addgene, Watertown, MA). For AAV barcode library production, we used pdsAAV-CAG-VBCx plasmids (where VBC is viral barcode and x is an integer identification number indicating each different viral barcode contained in each plasmid). pdsAAV-CAG-VBCx plasmids are double-stranded AAV vector plasmids and same as pdsAAV-U6-VBCx described previously^53^ except that human U6 small nuclear RNA promoter has been replaced by the CAG promoter and SV40 polyadenylation signal has been incorporated (Nakai, H., Huang, S., Adachi, K. US11,459,558. October 4, 2022.) .

### AAV Barcode-Seq analysis

Total DNA was extracted from tissues using QIAamp MinElute Virus Spin Kit (57704, Qiagen, Venlo, Netherlands) or KingFisher Cell and Tissue DNA kit (97030196, Fisher Scientific, Hampton, NH) following Proteinase K (25530049, Invitrogen, Waltham, MA) treatment. Total RNA was extracted from tissues using TRIzol (15596018, Invitrogen, Waltham, MA) followed by DNase treatment using TURBO DNA-free kit (AM1907, Invitrogen, Waltham, MA). Point eight μg of DNase-treated RNA was reverse-transcribed with a reverse transcription (RT)-specific primer using High-Capacity cDNA Reverse Transcription Kit (4368813, Applied Biosystems, Waltham, MA) or SuperScript IV Reverse Transcriptase (18090200, Invitrogen, Waltham, MA) in a total volume of 20 μL. One μg DNA or 4 μL cDNA was used to PCR-amplify virus barcode (VBC) using Platinum SuperFi II DNA Polymerase (12361010, Invitrogen, Waltham, MA). Information on primer sequence including frameshifting nucleotide, sample barcode was previously explained.^17, 46^ PCR products were mixed at an equimolar ratio and sequenced with either of the following settings at Massively Parallel Sequencing Shared Resource (MPSSR) at OHSU or Novogene (Sacramento, CA). The sequencing was performed using the following configurations on an Illumina NextSeq 500 or NovaSeq 6000 instrument: 75-cycle single-end, 150-cycle single-end, 180-cycle single-end and 300-cycle paired-end. We analyzed the Illumina sequencing data at the Pittsburgh Supercomputing Center or the Advanced Computing Center at OHSU. Four parameters in FastQC (per base sequence quality, per sequence quality scores, per base N content and sequence length distribution) were met in all the data sets we used in this study. The algorithm for data analysis was previously described.^20, 53^ For RNA Barcode-seq, the sequence-dependent differences in the *in vivo* mRNA transcription and the reverse-transcription (RT) PCR amplification efficiencies between VBCs were considered. In brief, we made two AAV9-CAG-VBCx libraries that contain all the AAV-CAG-VBCx genomes packaged with the same AAV9 capsid. These two libraries were produced independently from two independent pools of all the pdsAAV-CAG-VBCx plasmids mixed at an equimolar ratio. Each of the two AAV9-CAG-VBCx virus libraries was intravenously injected into 8-week-old C57BL/6J male mice (n=3 each) to obtain an independent, duplicated set of data each of which was obtained from 3 mice. The livers were harvested from the library-injected mice 6 weeks post-injection. Liver RNA and DNA was extracted and then subjected to the Barcode-Seq analysis, which provides RNA and DNA barcode reads, respectively. The correction factors obtained by the ratio of RNA and DNA barcode reads were used to cancel out the barcode sequence-dependent differences in the *in vivo* mRNA transcription and the RT-PCR amplification efficiencies between VBCs.

### AAV vector genome quantification

Total DNA was extracted from tissues as described in the AAV Barcode-Seq analysis section. Blood DNA sample was prepared using Extract-N-Amp Blood PCR Kit (XNAB2R, Sigma, St. Louis, MO) and diluted 100 times. Total DNA was extracted from 100 μL urine using Wako DNA Extraction Kit (29550201, Wako Chemicals, Richmond, VA) following Proteinase K treatment. Urine DNA pellet was dissolved in 15 μL Tris-HCl buffer. AAV vector genome copy numbers were quantified by quantitative PCR (qPCR). In brief, 10-100 ng of tissue DNA or 1 μL of blood or urine DNA sample was mixed with Power SYBR Green PCR Master Mix (43-676-59, Fisher Scientific, Hampton, NH) and 25 pmol primers in a 25 μL reaction volume and subjected to qPCR using Rotor-Gene Q (Qiagen, Venlo, Netherlands). We amplified the tdTomato gene sequence for vector genome quantification using the following primers: tdTomato forward (5’- ATGGACCTGTGATGCAGAAG-3’) and tdTomato reverse (5’- TTCAGCTTCAGAGCCTGGTG-3’). Information on copy number standards and normalization was described previously^20^. Vector genome copy numbers were expressed as double-stranded vector genome copy numbers per diploid genomic equivalent (ds-vg/dge).

### Immunofluorescence microscopy

Organs were harvested from mice following PBS perfusion. Harvested organs were fixed in 4% paraformaldehyde (PFA) (158127, Sigma, St. Louis, MO) and subsequently equilibrated in 30% sucrose (S5, Fisher Scientific, Hampton, NH) for cryoprotection. The fixed tissues were cryo-embedded in Tissue-Tek O.C.T. Compound (4583, Sakura Finetek, St. Torrance, CA) and cut into 5-μm-thick sections using a cryostat. Staining was performed as described previously^54^ using rabbit anti-WT1 (1:100, ab89901, Abcam, Cambridge, UK), rat anti-CD31 (1:100, 550274, BD Biosciences, Franklin Lakes, NJ), Lotus Tetragonolobus Lectin (LTL)-fluorescein (1:400, FL-1321-2, Vector Laboratories, Newark, CA), goat anti-rabbit IgG Alexa Fluor 647 (1:500, 111-605-144, Jackson ImmunoResearch, West Grove, PA), goat anti-rat IgG Alexa Fluor 647 (1:500, 112-605-167, Jackson ImmunoResearch, West Grove, PA) and Hoechst (1:10000, H3570, ThermoFisher Scientific, Waltham, MA). Immunofluorescence images were obtained using an BZ-X700 Fluorescence Microscope (Keyence, Osaka, Japan) or an LSM900 Confocal Microscope with Airyscan2 (Carl Zeiss, Oberkochen, Germany). We quantified the percentage of tdTomato-positive area in LTL-positive proximal tubules using Fiji version of ImageJ (NIH).^55^ At least 4 images per kidney were analyzed. To determine the percentage of tdTomato-positive podocytes, five glomeruli per kidney were selected from sections that were cut in close proximity to the middle of the glomeruli. Z-stacked images were taken and tdTomato-positive and -negative podocytes were manually counted. For the quantitative analysis of heart transduction, the heart area was determined through thresholding, and the mean fluorescent intensities (MFIs) of both the heart and the background was measured by ImageJ. The net intensity obtained by subtracting the background MFI from the heart MFI was used for the quantification of transduction.

### Assessment of nanoparticle distribution in the kidney following injection

Eight-week-old C57BL/6J male mice were injected with fluorescent polystyrene beads (G25, ThermoFisher Scientific, Waltham, MA)^56^ via IV (300 μL), RV (50 μL) and RP (50 μL). The beads are fluorescent microspheres with a diameter of 25 nm, which is comparable in size to AAV vector particles. Therefore, they can be used as surrogates to investigate the distribution of AAV particles following injection. We used 4 conditions, IV, RV, RP with blockage of renal artery and vein and RP without blockage of renal artery and vein (n=1 per condition). For IV injection, the distribution was assessed 30 min post-injection. For RV and RP injections, the distribution was assessed immediately after the completion of each injection procedure. Organs were harvested after PBS perfusion to remove beads from the blood vessels. Staining was performed as described above.

### Assessment of albuminuria

Spot urine was collected for the quantification of albuminuria. Urine albumin concentration was measured using Albuwell M (1011, Ethos Biosciences, Logan Township, NJ) and normalized by creatinine concentration measured by Creatinine Reagent Assay (C75391250, Pointe Scientific, Canton, MI). The data was expressed as gram per gram creatinine.

## Blinding

The following procedures were conducted in a blinded fashion in which the investigator remained unaware of the specifics of the vectors and samples: AAV Barcode-Seq analyses, urine albumin measurement, and a part of fluorescence microscopic studies in healthy and CKD kidneys. Although other experiments were conducted in an unblinded fashion, efforts were made to minimize bias and ensure objective observations.

### Statistics

Statistical analyses were performed using GraphPad Prism9 (version 9.5.0). Data are presented as mean ± standard error of the mean. Due to the small sample sizes in our study, normality tests such as the Shapiro Wilk test may have limited power in detecting departures from normality. Therefore, we assumed approximate normality based on the visual inspection of the data and conducted parametric tests, the results of which are presented in this paper. In this approach, comparisons between two groups were performed using a two-tailed Welch’s t-test. For data sets with more than two groups, significance was determined by a one-way ANOVA or two-way ANOVA followed by Tukey’s post-hoc test, or a two-tailed Welch’s t-test with Bonferroni correction. For pharmacokinetic study between AAV9 and AAV-KP1, significance was determined by a two-way repeated ANOVA followed by Bonferroni’s post-hoc test. P<0.05 was considered as statistical significance.

## Data availability

The NGS datasets for capsid libraries reported in this article are available under Sequence Read Archive accession code PRJNA998389 (http://www.ncbi.nlm.nih.gov/bioproject/998389). The data that support the findings of this study are available from the corresponding author upon reasonable request.

## Code availability

The scripts used for the Barcode-Seq analysis have been deposited into GitHub and can be accessed via https://github.com/nakaih-ohsu/AAV_Kidney_2023.git.

## Acknowledgements

The work involving RP injection was supported in part by a Sponsored Research Fund from Otsuka Pharmaceutical Co., Ltd. This work used the Extreme Science and Engineering Discovery Environment (XSEDE), which is supported by National Science Foundation grant number ACI-1548562. Specifically, it used the Bridges-2 system, which is supported by NSF award number ACI-1928147, at the Pittsburgh Supercomputing Center (PSC). We thank Guangping Gao and James M. Wilson for AAV helper plasmids including those for AAV8 and AAV9, Katja Pekrun and Mark A. Kay for helper plasmids for AAV-KP1, AAV-KP2 and AAV-KP3, Frank Park for the advice for the project and the manuscript, Yujiro Maeoka and James A. McCormick for the assistance of metabolic cage experiment, Helen R. Baggett for AAV vector production, Colton Stensrud for the assistance of mice breeding, Samuel J. Huang for the advice for fluorescence microscopy, and Kazuhiro Takahama and Yoshinori Katayama for their assistance of renal pelvis injection and pharmacokinetic study, respectively.

## Author contributions

T.F. and H.N. designed the study and wrote the manuscript. K.A. produced the AAV library and characterized AAV vectors following intravenous administration. T.F. and M.G.L. performed fluorescence microscopy and urine albumin ELISA. H.N. developed the algorithm for data analysis and wrote the computer scripts. Other experiments were performed by T.F. A.S and R.D. critically reviewed the paper. All authors read and approved the final version of the paper.

## Declaration of interests

H.N. receives royalty of AAV-related technologies licensed by Takara Bio Inc. and Capsigen Inc., and serves as a consultant for biotech companies, and is a co-founder of Capsigen Inc., and hold shares of Capsigen Inc. and Sphere Gene Therapeutics.

## Declaration of Generative AI and AI-assisted technologies in the writing process

During the preparation of this work the authors used ChatGPT for proofreading assistance while acknowledging its limitations. After using the tool, the authors reviewed and edited the content as needed and take full responsibility for the content of the publication.

## Legends for Supplemental Figures

**Figure S1.**
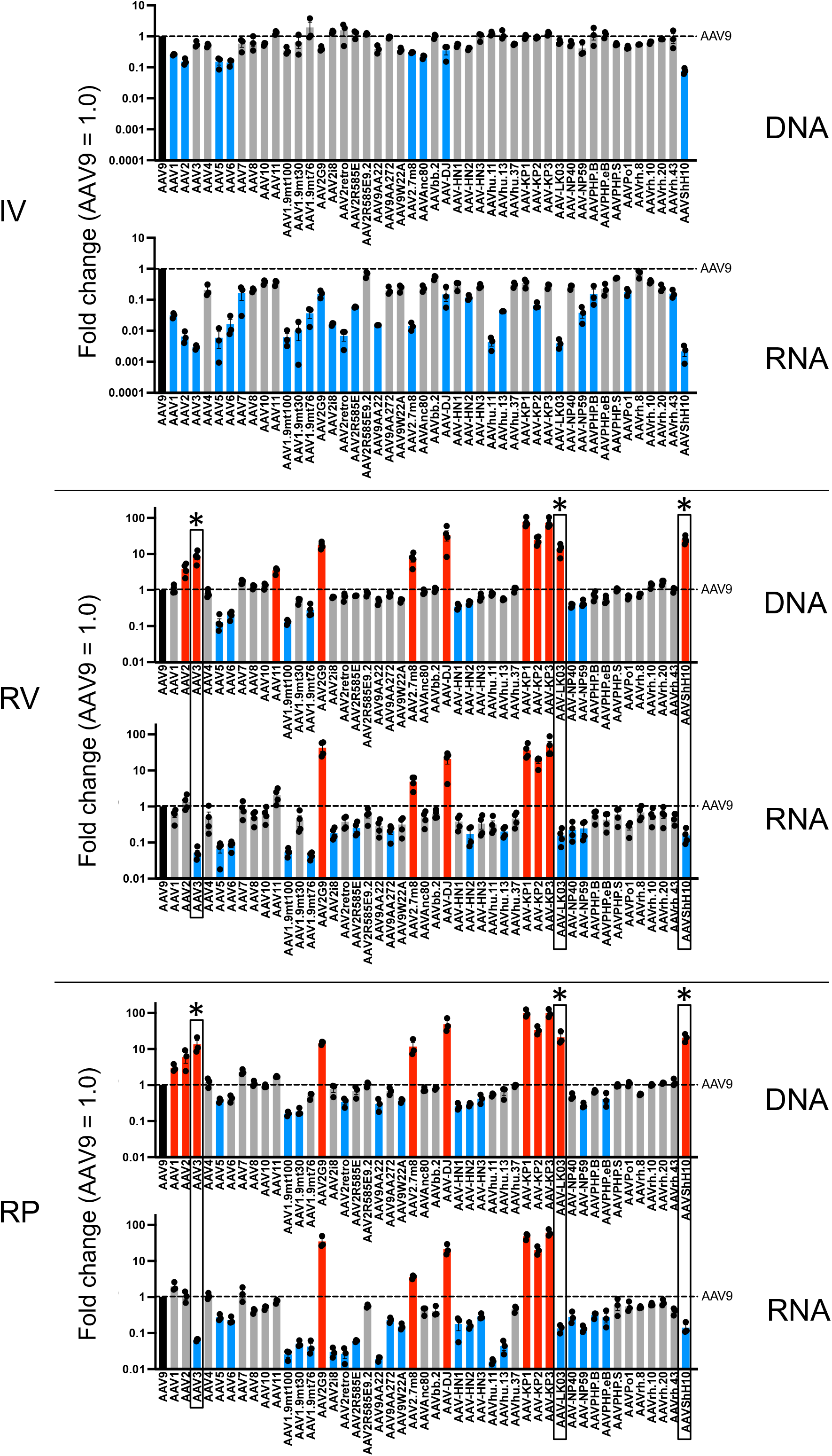
AAV Barcode-Seq analysis of AAV capsids for renal transduction following intravenous (IV), renal vein (RV) and renal pelvis (RP) injections. Eight-week-old C57BL/6J male mice were injected with an AAV-CAG-Barcode library that contained 47 AAV capsids via IV (2x10^13^ vg/kg), RV (3x10^11^ vg/mouse) or RP (3x10^11^ vg/mouse) (n=3 for IV and RP, n=4 for RV). Kidneys were harvested 6 weeks post-injection for the AAV Barcode-Seq analysis. Relative quantities of each AAV capsid-derived vector genomes (DNA) and vector genome transcripts (RNA) in the kidney were determined by AAV Barcode-Seq and expressed as fold changes. AAV capsids that showed significantly higher and lower renal transduction than AAV9 were highlighted with red and blue, respectively. AAV capsids that showed significantly higher renal transduction than AAV9 at RNA level by RV or RP injection were included in Figure 1. The AAV capsids indicated with an asterisk (*) are those that exhibited significantly enhanced DNA delivery to the kidney while mediating transgene expression at significantly lower levels compared to AAV9. Error bars indicate SEM. A one-way ANOVA followed by Tukey’s post hoc test was used for statistical assessment of the data. Adjusted p<0.05 was considered as statistical significance.

**Figure S2.**
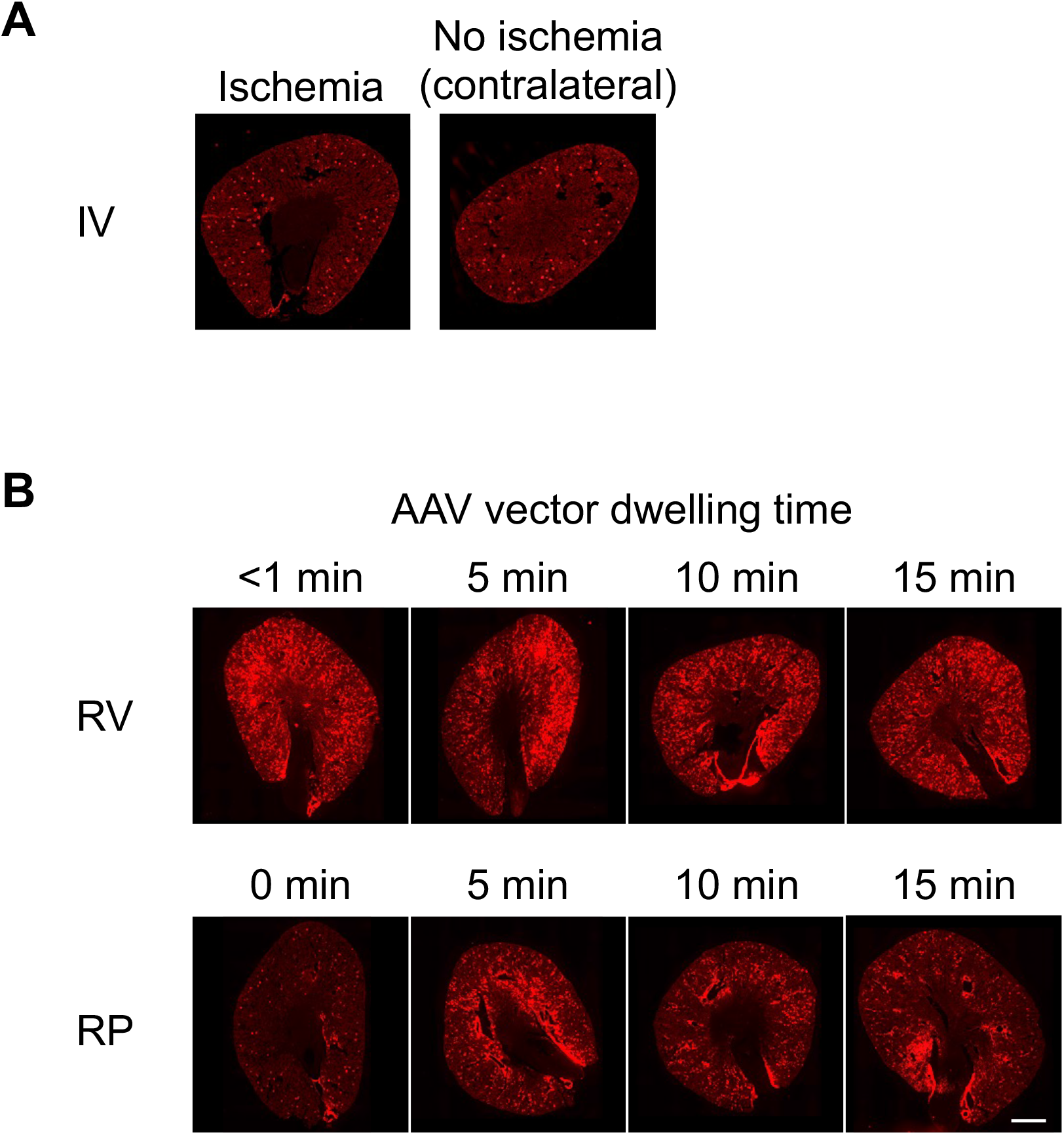
Effects of ischemia and AAV vector dwelling time on renal transduction. Eight-week-old C57BL/6J male mice were injected with AAV-KP1-CAG-tdTomato at a dose of 3x10^11^ vg/mouse. Kidneys were harvested 2 weeks post injection for downstream analyses. Representative fluorescence microscopic images in each condition are shown. (A) The AAV-KP1 vector was administered intravenously with or without a 15-min renal ischemic time prior to injection (n=1). **(B)** The AAV-KP1 vector was administered via RV or RP injection with different durations of AAV vector dwelling time (n=1 each). Note that renal blood flow needs to be stopped for less than 1 min to perform RV injection. Scale bar, 1 mm.

**Figure S3.**
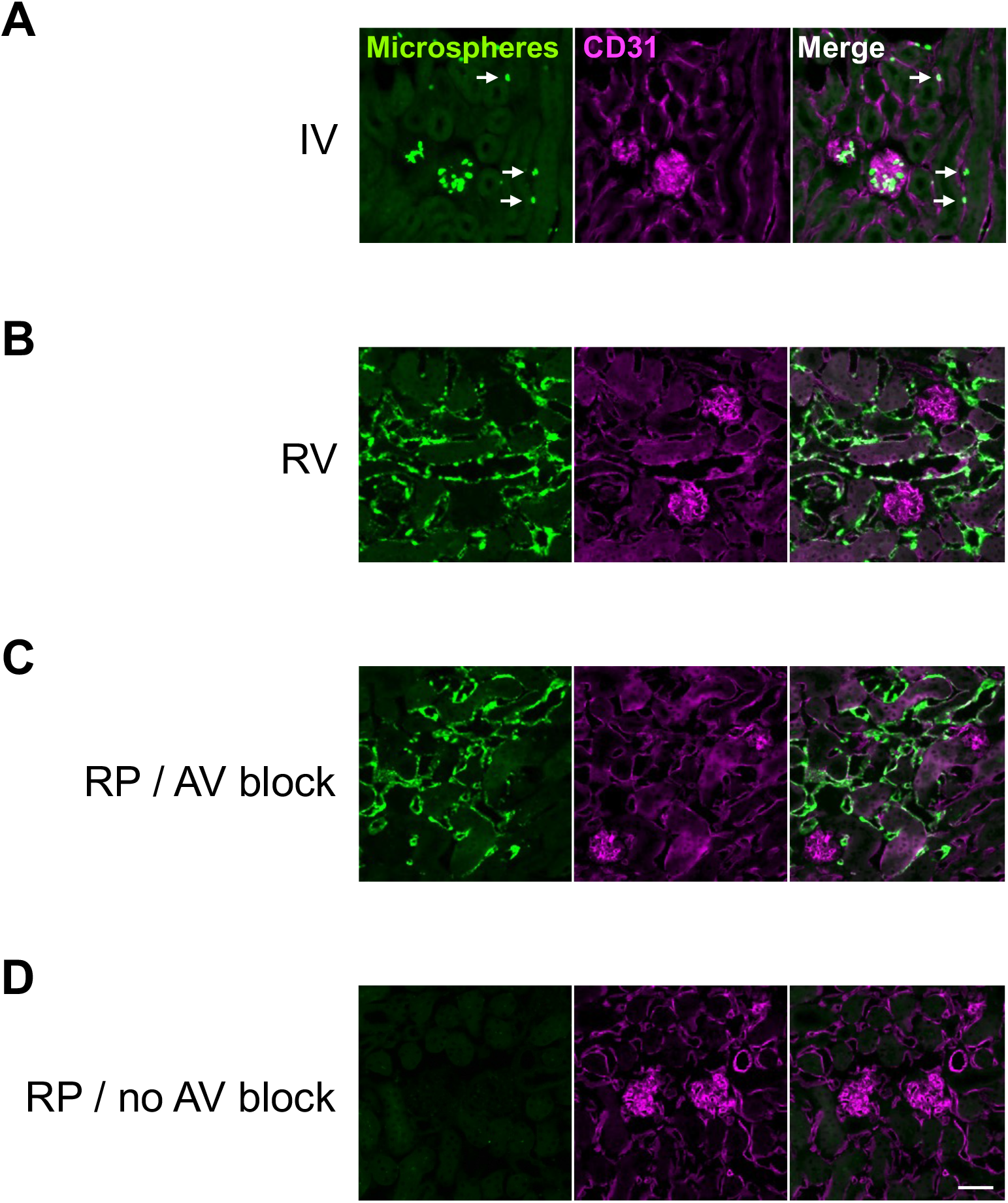
Intrarenal distribution of 25 nm microspheres following IV, RV and RP injections. Eight-week-old C57BL/6J male mice were injected with fluorescent 25 nm microspheres via IV (A), RV (B) and RP with (C) or without (D) blockage of the renal blood flow (n=1 each). For the IV injection, the microspheres were circulated in the body for 30 min. There was no circulation time for RV and RP injections. Kidneys were harvested following PBS perfusion immediately after the completion of each injection procedure. Representative fluorescence microscopic images of the treated kidneys are shown. White arrows in Panel A show accumulation of microspheres in the renal interstitium. Green, microspheres; magenta, CD31 (a marker for endothelial cells). Scale bar, 50 μm.

**Figure S4.**
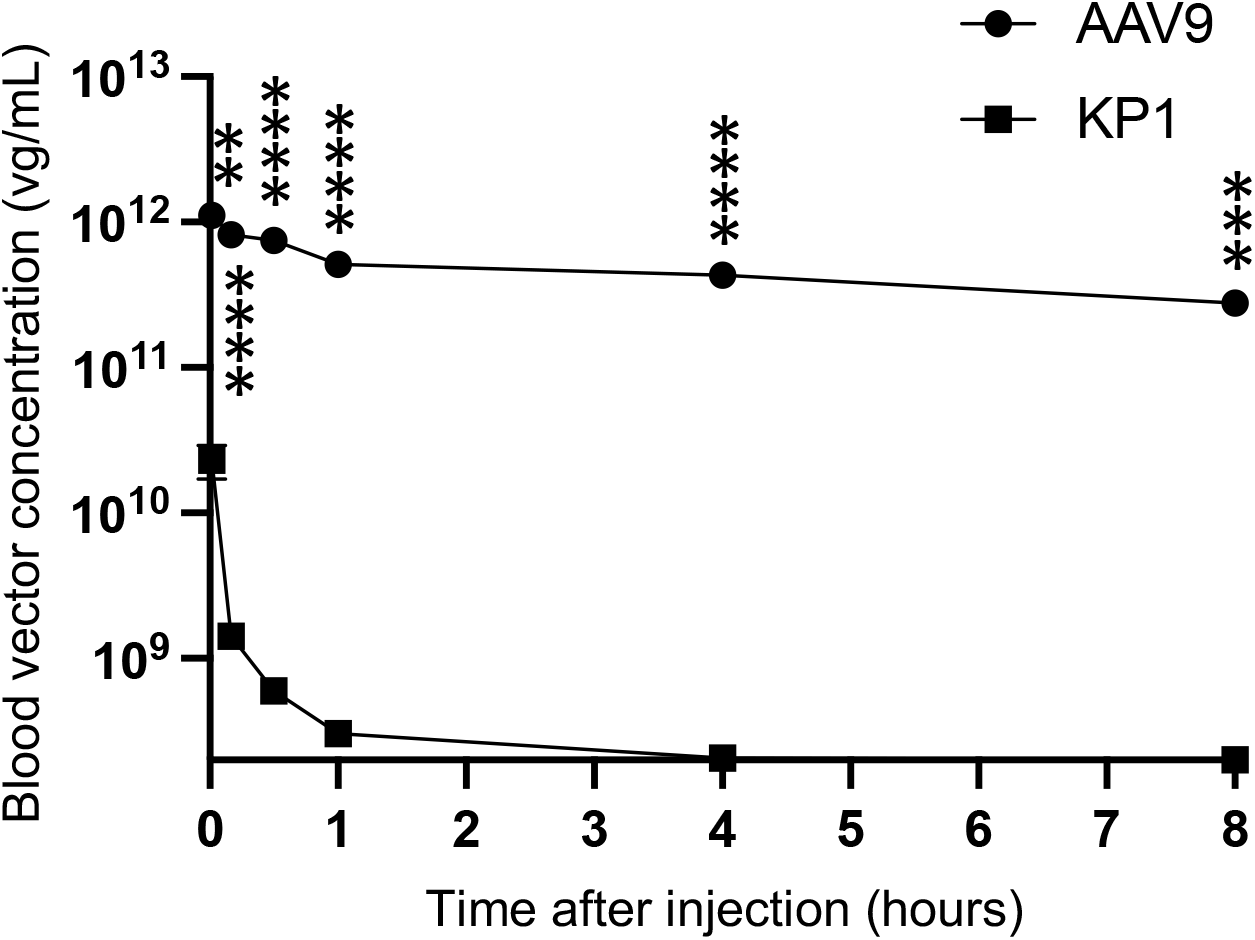
Pharmacokinetics of AAV9 and AAV-KP1 in the blood following IV injection. Eight-week-old C57BL/6J male mice were injected via IV with AAV9-CAG-tdTomato and AAVKP1-CAG-tdTomato at a dose of 1x10^13^ vg/kg (n=4 per group). A two-way repeated ANOVA followed by Bonferroni’s post hoc test was used for statistical assessment of the data. The p value represents the statistical significance between AAV9 and AAV-KP1 at each time point. **, adjusted p<0.01; ***, adjusted p<0.001; ****, adjusted p<0.0001. Bars represent SEM.

**Figure S5.**
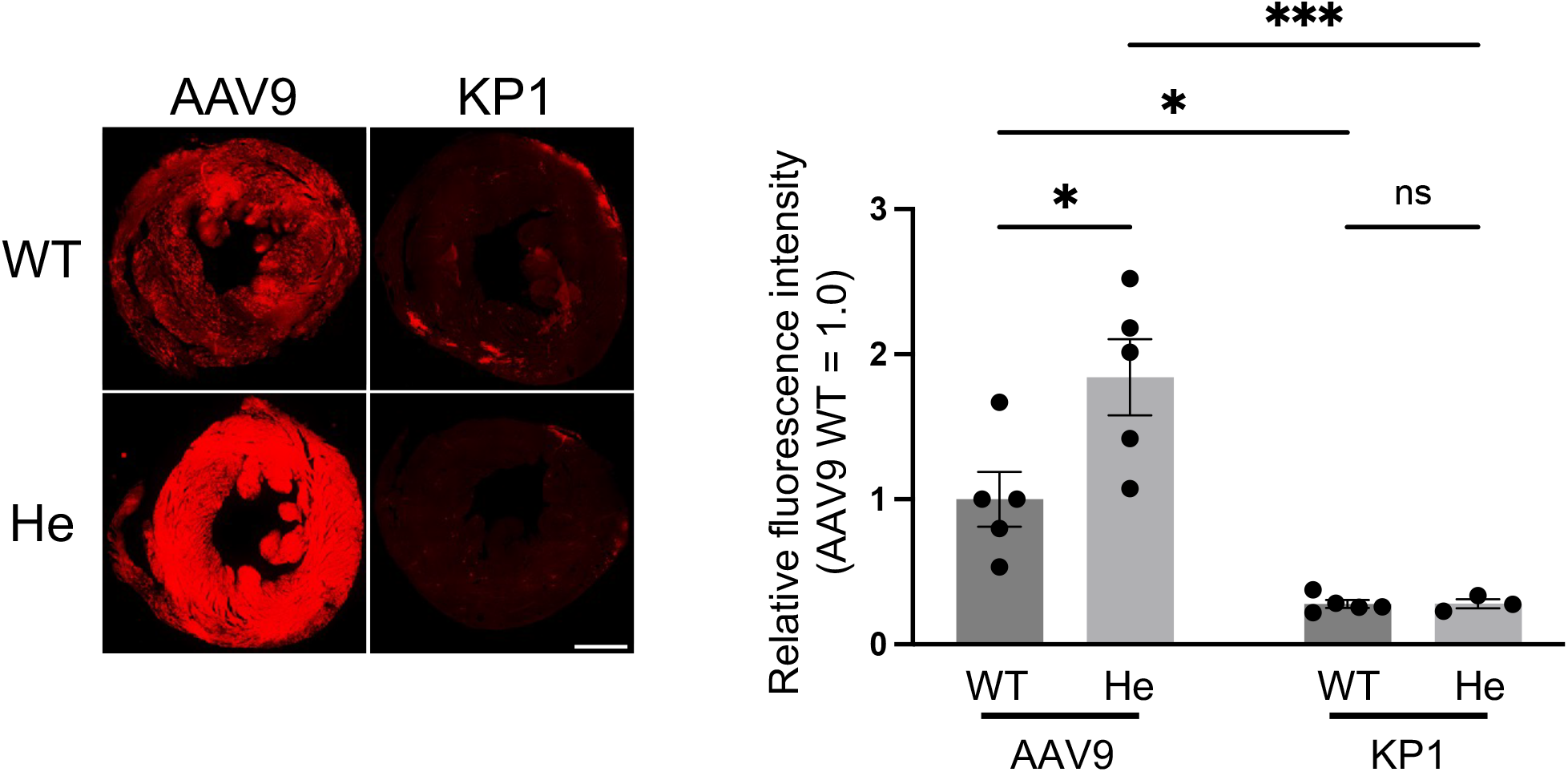
Enhanced cardiac transduction with AAV9 in CKD. Twenty-five to 30-week-old Col4a5 G5X hemizygous male mice (He) and age-matched wild-type control male mice (WT) were injected with AAV9-CAG-tdTomato and AAV-KP1-CAG-tdTomato via IV at a dose of 1x10^13^ vg/kg. Hearts were harvested 2 weeks post-injection for downstream analyses. AAV9 in WT (n=5), AAV9 in He (n=5), AAV-KP1 in WT (n=5) and AAV-KP1 in He (n=3). **(A)** Representative fluorescence microscopic images of native tdTomato in the heart in the AAV vector-treated mice. **(B)** AAV9 and AAV-KP1 transduction efficiencies in the heart. Cardiac transduction efficiency relative to that of AAV9 in WT was assessed by quantifying the mean fluorescence intensity of tdTomato in the heart. Fiji version of ImageJ was used for the image analysis. A two-way ANOVA followed by Tukey’s post hoc test was used for statistical assessment of the data. *, adjusted p<0.05; ***, adjusted p<0.001; ns, not significant. Scale bars, 1 mm. Error bars represent SEM.

**Table S1.** AAV capsids included in the Barcode library.

(<sup>a</sup> described in Nakai, H., Huang, S., Adachi, K. US11,459,558. October 4, 2022.)
| **Capsids** | **Reference** | **Capsids** | **Reference** |
| --- | --- | --- | --- |
| AAV1 | Earley<sup>53</sup> | AAVbb.2 | Gao<sup>57</sup> |
| AAV1.9mt100 | Our lab | AAV-DJ | Grimm<sup>22</sup> |
| AAV1.9mt30 | Our lab | AAV-HN1 | Our lab<sup>a</sup> |
| AAV1.9mt76 | Our lab | AAV-HN2 | Our lab<sup>a</sup> |
| AAV2 | Earley<sup>53</sup> | AAV-HN3 | Our lab<sup>a</sup> |
| AAV2G9 | Shen<sup>23</sup> | AAVhu.11 | Gao<sup>57</sup> |
| AAV2i8 | Asokan<sup>58</sup> | AAVhu.13 | Gao<sup>57</sup> |
| AAV2retro | Tervo<sup>59</sup> | AAVhu.37 | Gao<sup>57</sup> |
| AAV2R585E | Adachi<sup>20</sup> | AAV-KP1 | Pekrun<sup>21</sup> |
| AAV2R585E9.2 | Adachi<sup>20</sup> | AAV-KP2 | Pekrun<sup>21</sup> |
| AAV3 | Earley<sup>53</sup> | AAV-KP3 | Pekrun<sup>21</sup> |
| AAV4 | Earley<sup>53</sup> | AAV-LK03 | Lisowski<sup>60</sup> |
| AAV5 | Earley<sup>53</sup> | AAV-NP40 | Paulk<sup>61</sup> |
| AAV6 | Earley<sup>53</sup> | AAV-NP59 | Paulk<sup>61</sup> |
| AAV7 | Earley<sup>53</sup> | AAVPHP.B | Deverman<sup>62</sup> |
| AAV8 | Earley<sup>53</sup> | AAVPHP.eB | Chan<sup>63</sup> |
| AAV9 | Earley<sup>53</sup> | AAVPHP.S | Chan<sup>63</sup> |
| AAV9AA272 | Our lab | AAVpo1 | Bello<sup>64</sup> |
| AAV9AA22 | Our lab | AAVrh.8 | Gao<sup>57</sup> |
| AAV9W22A | Our lab | AAVrh.10 | Gao<sup>57</sup> |
| AAV10 | Earley<sup>53</sup> | AAVrh.20 | Gao<sup>57</sup> |
| AAV11 | Earley<sup>53</sup> | AAVrh.43 | Gao<sup>57</sup> |
| AAV2.7m8 | Dalkara<sup>24</sup> | AAVShH10 | Klimczak<sup>65</sup> |
| AAVAnc80 | Zinn<sup>66</sup> |  |  |

## Notes

### Competing Interest Statement

H.N. receives royalty of AAV-related technologies licensed by Takara Bio Inc. and Capsigen Inc., serves as a consultant for biotech companies, and is a co-founder of Capsigen Inc., a company commercializing AAV vector-related technologies.

